# Stochastic chain termination in bacterial pilus assembly

**DOI:** 10.1101/2022.10.11.511808

**Authors:** Christoph Giese, Chasper Puorger, Oleksandr Ignatov, Zuzana Bečárová, Marco E. Weber, Martin A. Schärer, Guido Capitani, Rudi Glockshuber

**Affiliations:** Institute of Molecular Biology and Biophysics, Department of Biology, ETH Zurich, 8093 Zurich, Switzerland; Present address: Institute for Chemistry and Bioanalytics, University of Applied Sciences and Arts Northwestern Switzerland, 4132 Muttenz, Switzerland; Present address: V.I. Grishchenko Clinic of Reproductive Medicine, Blahovishchenska st.25, 61052 Kharkiv, Ukraine; Present address: Laboratory of Physical Chemistry, Department of Chemistry and Applied Biosciences, ETH Zurich, 8093 Zurich, Switzerland; Laboratory of Biomolecular Research, Paul Scherrer Institute, 5232 Villigen-PSI, Switzerland

**Keywords:** Uropathogenic *E. coli*, catalysis of protein assembly, type 1 pili, assembly termination, FimI, assembly kinetics, filamentous protein polymers

## Abstract

Adhesive type 1 pili from uropathogenic *Escherichia coli* strains are filamentous, supramolecular protein complexes consisting of a short tip fibrillum and a long, helical rod formed by up to several thousand copies of the major pilus subunit FimA. Here, we reconstituted the entire type 1 pilus rod assembly reaction *in vitro*, using all constituent protein subunits in the presence of the assembly platform FimD, and identified the so-far uncharacterized subunit FimI as an irreversible assembly terminator. We provide a complete, quantitative model of pilus rod assembly kinetics based on the measured rate constants of FimD-catalyzed subunit incorporation. The model reliably predicts the length distribution of assembled pilus rods as a function of the ratio between FimI and the main pilus subunit FimA and is fully consistent with the length distribution of membrane-anchored pili assembled *in vivo*. The results show that the natural length distribution of adhesive pili formed via the chaperone-usher pathway results from a stochastic chain termination reaction.

## INTRODUCTION

Many Gram-negative pathogens use adhesive, filamentous protein complexes anchored to their outer membrane, termed pili, to attach to surface glycans of host cells and initiate infection (Connell et al., 1996; Jones et al., 1995; Lindberg et al., 1987; Martinez et al., 2000; Roberts et al., 1994). Type 1 pili and the related P pili belong to the best-characterized pilus systems from uropathogenic *Escherichia coli* strains (Hospenthal and Waksman, 2019). Type 1 pili bear a single copy of the lectin (adhesin) FimH at their distal end, which recognizes terminal mannosides in high-mannose type N-glycans of the urothelial receptor uroplakin Ia (Xie et al., 2006; Zhou et al., 2001). FimH is bound to one or several copies of the minor subunits FimG and FimF (Hahn et al., 2002; Jones et al., 1995; Le Trong et al., 2010), which, together with FimH, form a short, linear tip fibrillum (Figure 1A). The tip fibrillum is connected to the pilus rod, a helical and rigid quaternary structure composed of hundreds to several thousand copies of the main structural pilus subunit FimA (Brinton, 1965; Hospenthal et al., 2017). Pilus assembly *in vivo* follows the chaperone-usher pathway (Hospenthal and Waksman, 2019; Werneburg and Thanassi, 2018) and is strictly dependent on two protein catalysts: the periplasmic chaperone FimC that accelerates pilus subunit folding in the periplasm up to 10^4^-fold, and the assembly platform (“usher”) FimD in the outer membrane that catalyzes pilus subunit polymerization independently of ATP (Crespo et al., 2012; Jacob-Dubuisson et al., 1994; Klemm, 1992; Klemm and Christiansen, 1990; Nishiyama et al., 2008; Vetsch et al., 2004).

**Figure 1.**
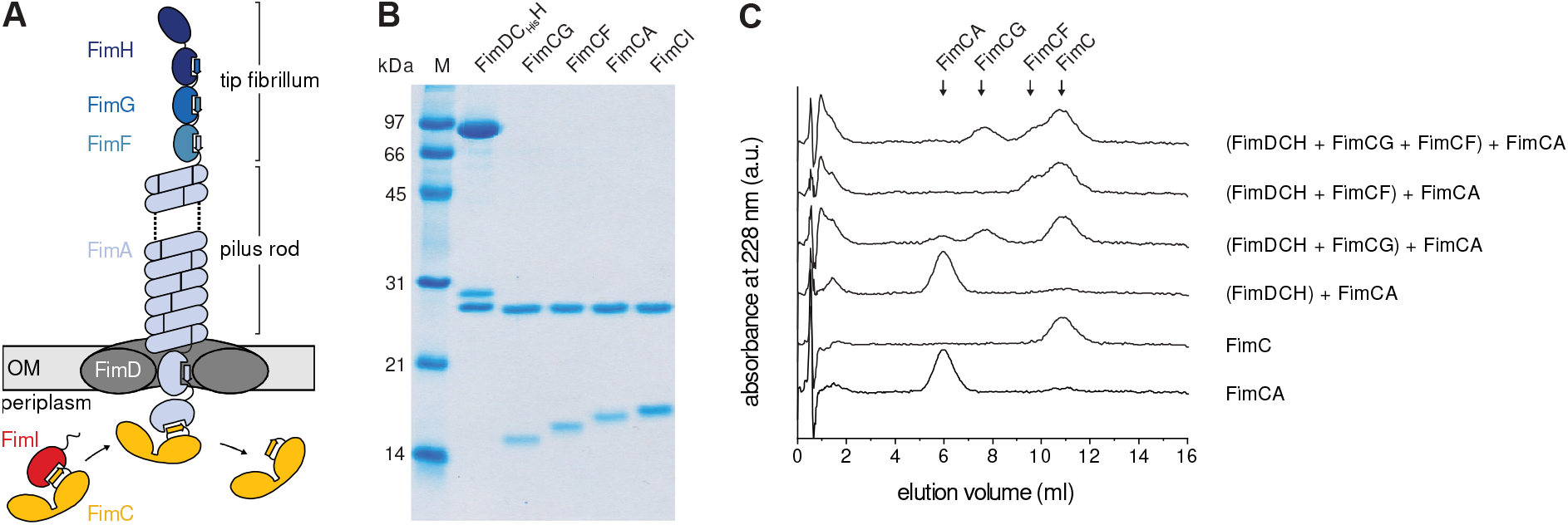
*In vitro* reconstitution of type 1 pilus rod assembly. (A) Architecture and subunit composition of the type 1 pilus. The linear tip fibrillum consists of the adhesin FimH and the minor subunits FimG and FimF. It is attached to the helical pilus rod that may consist of several thousand copies of the main subunit FimA (Brinton, 1965; Hahn et al., 2002; Jones et al., 1995). Pilus assembly is catalyzed by the assembly platform FimD in the outer membrane (OM) (Nishiyama et al., 2008). Only subunits bound to the periplasmic chaperone FimC are assembly-competent and recognized by FimD (Nishiyama et al., 2005; Nishiyama et al., 2003). The mechanism of incorporation of the assembly terminator subunit FimI (red) is the focus of this study. (B) Coomassie-stained polyacrylamide-SDS gel of the purified type 1 pilus subcomplexes (25 pmol each) used in this study. (C) Analytical cation ion exchange chromatography runs, monitoring the decrease in the concentrations of free FimCA complexes during FimD-catalyzed pilus rod assembly *in vitro*. FimDCH (0.35 μM) was preincubated with or without an 8-fold molar excess of FimCG or FimCF or both for 5 min at 37 °C. After addition of FimCA (final concentrations of FimDCH and FimCA were 0.25 and 5 μM, respectively) and incubation for 10 min at 37 °C, the reactions were stopped by rapid cooling on ice. The remaining, free FimC-subunit complexes were separated and quantified by cation exchange chromatography at 4 °C and pH 6.7. Reference elution profiles for 5 μM FimC and 5 μM FimCA are shown at the bottom of the panel.

The adhesin FimH is a two-domain protein with an N-terminal lectin domain that binds glycoprotein receptors, and a C-terminal pilin domain that links FimH to the next subunit FimG (Choudhury et al., 1999; Hahn et al., 2002; Le Trong et al., 2010; Xie et al., 2006; Zhou et al., 2001). In contrast to FimH, all other pilus subunits are single-domain proteins and structural homologs of the FimH pilin domain, characterized by an incomplete immunoglobulin (Ig)-like fold lacking the C-terminal β-strand (Choudhury et al., 1999). After secretion of the unfolded subunits into the periplasm via the SecYEG system, the oxidoreductase DsbA catalyzes formation of an invariant structural disulfide bond connecting β-strands A and B in each of the subunits (Crespo et al., 2012). The periplasmic chaperone FimC then specifically recognizes the disulfide forms of the unfolded subunits and catalyzes subunit folding (Crespo et al., 2012; Vetsch et al., 2004). In all native FimC-subunit complexes, FimC completes the Ig-like subunit fold via donor strand complementation (DSC) by inserting its β-strand G1 into the incomplete Ig fold of the subunit in a parallel orientation relative to the subunit’s C-terminal β-strand F (Choudhury et al., 1999; Crespo et al., 2012; Eidam et al., 2008). Except for the distal subunit FimH, all other subunits possess an N-terminal extension (Nte, also termed donor strand) of approx. 15‒20 amino acids (Choudhury et al., 1999) that remains exposed and unstructured in FimC-subunit complexes.

Only native FimC-subunit complexes are assembly-competent and specifically recognized by the assembly catalyst FimD in the outer membrane (Nishiyama et al., 2005; Nishiyama et al., 2003). FimD consists of five domains: a central, 24-stranded β-barrel transmembrane domain that is obstructed by a plug domain in the inactive, resting state of FimD, an N-terminal and two C-terminal periplasmic domains (FimD_N_ and CTDs, respectively) (Du et al., 2021; Du et al., 2018; Geibel et al., 2013; Huang et al., 2009; Nishiyama et al., 2003; Omattage et al., 2018; Phan et al., 2011; Remaut et al., 2008). Chaperone-subunit complexes are first bound by FimD_N_ (Ng et al., 2004; Nishiyama et al., 2003). Subunit incorporation into the pilus then occurs in an irreversible reaction termed donor strand exchange (DSE), in which the exposed N-terminal donor strand of the incoming subunit, bound to FimC and FimD_N_, displaces the chaperone capping the last incorporated subunit at the FimD translocation pore (Choudhury et al., 1999; Geibel et al., 2013; Hospenthal et al., 2017; Le Trong et al., 2010; Nishiyama et al., 2008; Remaut et al., 2006; Sauer et al., 1999; Sauer et al., 2002; Vetsch et al., 2006; Zavialov et al., 2003). In contrast to FimC-subunit complexes, the donor strand of the incoming subunit inserts in the opposite (antiparallel) orientation, which leads to extremely stable subunit-subunit interactions that practically show infinite stability against dissociation (Puorger et al., 2008). The donor strand acceptor groove of each subunit possesses five binding pockets, P1‒P5, that specifically accommodate hydrophobic side chains of the respective donor strand (Gossert et al., 2008; Hospenthal et al., 2017; Puorger et al., 2008; Remaut et al., 2006; Sauer et al., 2002). Structural data suggest that the DSE reaction is initiated by threading the donor strand of the incoming subunit into the P5 pocket of the acceptor subunit, which is not occupied when the acceptor subunit is capped with FimC (Remaut et al., 2006). During DSE, the incoming chaperone-subunit complex is handed over from FimD_N_ to the CTDs of FimD, thus resetting FimD_N_ for binding the next chaperone-subunit complex (Du et al., 2021; Du et al., 2018; Geibel et al., 2013; Omattage et al., 2018; Phan et al., 2011).

In the case of P pili, it is assumed that assembly termination is achieved when a sixth structural subunit, PapH, is incorporated into the pilus (Baga et al., 1987; Verger et al., 2006). The inhibitory action of PapH is attributed to the observation that the PapH structure, when complexed with the P pilus chaperone PapD, lacks a P5 pocket, which likely prevents donor strand attack and incorporation of another subunit via DSE (Verger et al., 2006). Besides its proposed function as assembly terminator, PapH also anchors P pili to the outer bacterial membrane (Baga et al., 1987). Despite these insights, termination of pilus assembly by irreversible incorporation of a terminator subunit has never been demonstrated directly for pili assembled via the chaperone-usher pathway, and the mechanism of type 1 pilus assembly termination has remained uncharacterized. Based on the similar gene arrangements in the P pilus and type 1 pilus gene clusters and the sequence similarity between the *papH* and *fimI* gene, FimI, the fifth structural subunit of type 1 pili, was suggested to act as functional equivalent of PapH (Rossolini et al., 1993). Here, we addressed this hypothesis with an *in vitro* study, in which we reconstituted the entire type 1 pilus assembly system from all purified components, including the previously uncharacterized subunit FimI. We demonstrate directly that FimI terminates type 1 pilus rod assembly and present a complete, quantitative kinetic description of pilus rod assembly and assembly termination. Specifically, the kinetics of subunit binding to FimD and irreversible subunit incorporation into the growing pilus are in full agreement with a stochastic chain termination mechanism and reliably predict length distribution histograms of pili formed *in vitro* and *in vivo* as a function of the FimI:FimA ratio. Unexpectedly, we find that incorporation of FimI is the fastest assembly step in pilus biogenesis, predicting a high excess of FimA over FimI during pilus assembly *in vivo*. Moreover, we show that incorporation of FimI is essential for stable anchoring of the pilus to the outer membrane. Finally, we present crystal structures of the FimC-FimI complex, the ternary FimD_N_-FimC-FimI complex and the ternary FimC-FimI-FimA complex. The latter represents the proximal end of the fully assembled pilus and provides a structural basis for understanding pilus assembly termination and pilus anchoring by FimI.

## RESULTS AND DISCUSSION

### Initiation of pilus rod assembly requires FimG or FimF at the proximal end of the tip fibrillum

To study the mechanism of the supposed type 1 pilus assembly terminator FimI, we reconstituted the entire FimD-catalyzed type 1 pilus assembly reaction *in vitro* from all purified components. To this end, we purified the ternary complex between FimD, FimC and FimH (FimDCH) and all binary complexes between the chaperone FimC and the pilus subunits FimG, FimF, FimA and FimI (FimCG, FimCF, FimCA and FimCI; Figure 1B). First, we probed the ability of FimDCH to catalyze pilus rod assembly from FimCA complexes by mixing FimDCH with a 20-fold excess of FimCA, using either FimDCH alone or FimDCH that had been preincubated with an 8-fold excess of FimCG or FimCF, or a FimCG/FimCF mixture. Kinetics of FimA polymerization were recorded by analytical cation exchange chromatography via the decrease in the concentration of free FimCA complexes and the increase in the concentration of free FimC released upon FimA polymerization (Figure 1C). While FimDCH alone proved to be inactive as catalyst of FimA assembly, complete turnover of FimCA and release of free FimC was observed when FimDCH had been preincubated with FimCG and/or FimCF prior to the addition of FimCA (Figure 1C).

Negative-stain electron microscopy confirmed that FimA polymers (pilus rods) had indeed formed when FimDCH had been pre-incubated with FimCG and/or FimCF (Figure S1). Thus, the initiation of FimD-catalyzed FimA assembly requires the presence of either a FimC-capped FimG or FimC-capped FimF subunit at the growing end of the pilus in the periplasm.

At first glance, the finding that FimDCH alone did not catalyze FimA assembly contradicts a previous study in which FimDCH catalyzed pilus rod assembly from FimCA complexes (Nishiyama et al., 2008). Very likely, tiny impurities with FimCG and FimCF in the previous preparation of FimCA are responsible for this discrepancy: Specifically, denatured and dissociated type 1 pili, i.e. a mixture of unfolded FimH, FimG, FimF and FimA, had served as source of FimA for production of FimCA complexes by refolding in the presence of FimC and purification of FimCA with ion exchange chromatography (Nishiyama et al., 2008). The resulting FimCA preparation thus may have contained trace amounts of FimCG and FimCF that allowed initiation of FimD-catalyzed pilus rod assembly (Nishiyama et al., 2008). In contrast, the FimCA complexes used in the present study were free of contaminations with FimCG and/or FimCF, as they were produced via refolding of FimA from cytoplasmically produced FimA aggregates (inclusion bodies; see Methods). In addition, the strict requirement of either FimF or FimG at the growing end of the tip fibrillum for initiation of pilus rod assembly (i.e., the incorporation of the first FimA copy after the tip fibrillum) found here fully agrees with the previous observation that FimD-catalyzed rod assembly was accelerated when purified FimCF and/or FimCG complexes were added (Nishiyama et al., 2008). The data also agree with studies on P pilus biogenesis and on type 1 pilus rod assembly in *Salmonella typhimurium*, where assembly of PapA to P pilus rods depended on the incorporation of pilus tip adaptor subunits PapF and PapK, and *S. typhimurium* type 1 pilus rod assembly required FimF (the only adaptor subunit present in *S. typhimurium*) (Jacob-Dubuisson et al., 1993; Zeiner et al., 2012). Together, the dependence of pilus rod assembly initiation on the presence of a minor tip fibrillum subunit appears to be common to all pilus systems with a heterooligomeric tip fibrillum. This mechanism, together with a very slow incorporation of the first FimA subunit (see below), also favors completion of the tip fibrillum prior to rod assembly. In contrast, formation of the chaperone-usher-adhesin complex alone appears to be sufficient for the assembly of pilus rods in pilus systems only bearing a single adhesin at the distal end of the pilus rod, such as pili from the Caf1, Afa/Dr or the CS1 family (Sakellaris and Scott, 1998; Zav’yalov et al., 2010).

We next analyzed the conditions required for converting FimDCH to the fully active FimA assembly catalyst. For this purpose, we pre-incubated FimDCH with FimCG, FimCF or both, then added excess FimCA, and measured the kinetics of FimD-catalyzed rod assembly as a function of pre-incubation time (Figure S2). Herein, we refer to this conversion as the “activation of FimDCH”, and restrict the term “catalytic activity” to the ability of FimD to catalyze FimA polymerization from FimCA and incorporation of the assembly terminator FimI (see below). When FimDCH was preincubated with FimCG alone, maximum FimD activity was attained after 30 min of preincubation. FimA assembly with 5 μM FimCA and 0.25 μM activated FimDCH took approximately 20 min for completion and FimD activity slowly decreased with longer preincubation times (Figure S2A). With FimCF alone, FimD activity was maximum only after 6 h of incubation, while FimCA assembly was completed in less than 15 min, indicating that FimD was a more efficient catalyst of pilus rod assembly when the first FimA subunit associated with FimF rather than FimG at the growing pilus end. In addition, no loss in FimD activity was observed for even longer preincubation times of up to 20 h (Figure S2B).

After initiation of FimD-catalyzed rod assembly by addition of excess (20-fold) FimCA to the activation reactions, all FimA assembly kinetics exhibited an initial lag-phase, indicating that the rate-limiting step corresponded to the incorporation of the first FimA subunit into the growing pilus (Nishiyama et al., 2008). We tested sequential (FimCG first, FimCF second) or simultaneous addition of FimCG and FimCF to FimDCH during the activation reaction and found the latter to yield the highest FimD activity, so that FimA assembly [0.18 μM activated FimDCH, 3.6 μM FimCA (20-fold excess)] was completed about 2-fold faster compared to sequential addition of FimCG and FimCF (Figure S2C). Pili generated with these most active FimD preparations were comparably short with a median length of only 0.6 · 10^2^ nm (Figure S2D). As expected, the increase of the FimCA:FimD ratio from 20:1 to 200:1 reproducibly yielded a median pilus length of 4.2 · 10^2^ nm (Figure S2E). In the following, we used these conditions for FimDCH activation as standard conditions to analyze the influence of the assembly terminator FimI on the lengths of pilus rods assembled via FimD.

### FimI terminates type 1 pilus assembly

To test whether FimI inhibited pilus rod assembly, purified FimCI (1 μM) was added to the FimD-catalyzed FimA assembly reaction (0.1 μM activated FimDCH, 20 μM FimCA) after 9 minutes, when rod assembly was completed to about 40% (Figure 2A). While the assembly reaction in absence of FimCI was completed within 20 minutes, the reaction stopped within about 5 minutes after FimCI addition and approximately half of FimCA remained unpolymerized, even after 65 minutes of incubation (Figure 2A). The length distributions of the formed pili (200 pili from each reaction) were analyzed by negative-stain electron microscopy. As expected for FimI functioning as the assembly inhibitor and in agreement with about 50% free FimCA complexes left after assembly termination by FimCI, the median pilus length decreased about 2-fold (from 3.8 · 10^2^ nm to 2.1 · 10^2^ nm) by FimCI, relative to the reaction in the absence of FimCI (Figure 2B).

**Figure 2.**
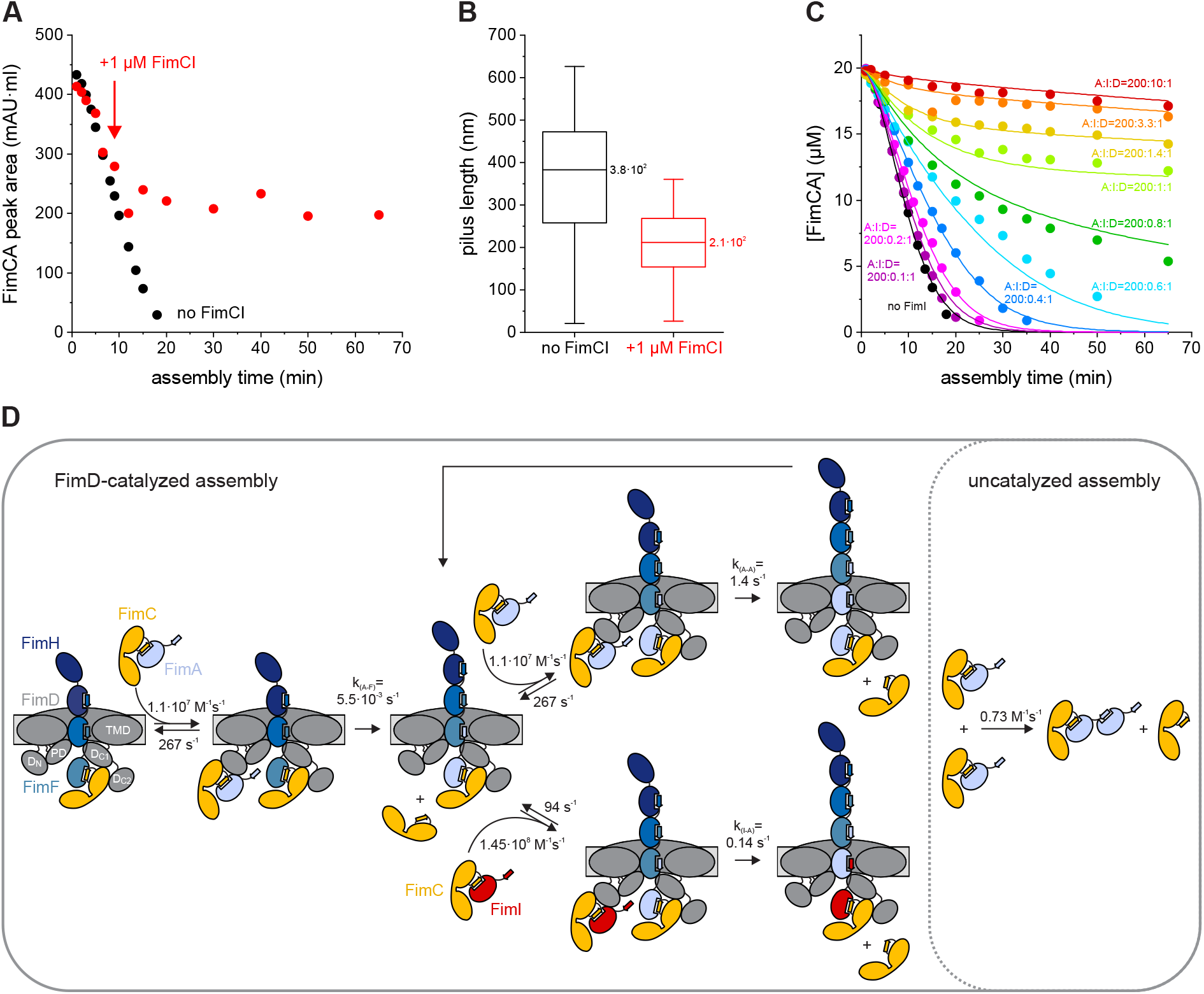
FimI inhibits type 1 pilus rod assembly. (A) Kinetics of FimD-catalyzed FimA assembly at pH 8.0 and 23 °C, recorded in absence of FimCI (black) or with FimCI added to 1 μM after nine minutes of the reaction (red). The time point of FimCI addition is indicated by an arrow. FimDCH (0.35 μM) was preincubated with an 8-fold molar excess of FimCG and FimCF for 30 min at 23 °C. After addition of FimCA (final concentrations of FimDCH and FimCA were 0.1 and 20 μM) and further incubation at 23 °C for defined periods of time, the reaction mixtures were analyzed by analytical cation exchange chromatography at 4 °C and pH 6.0. FimA assembly was monitored by recording the decrease in FimCA peak area with time. (B) Box plots of pilus length distributions of the two reactions shown in panel (A). Samples taken after 65 minutes of assembly were analyzed by negative-stain electron microscopy and the length of 200 pili each was measured. The box encloses the second and third quartile, the horizontal line indicates the median and the whiskers the smallest and largest value of the distribution (*n* = 200). (C) Kinetics of FimD-catalyzed FimA assembly in presence of different FimCI concentrations at pH 8.0 and 23 °C. Molar ratios between FimCA, FimCI and FimD (A:I:D) are indicated. Initial concentrations were 0.1 μM FimD, 20 μM FimCA and 0.01 – 1 μM FimCI. Solid lines show the result of a gobal fit of the data according to the model depicted in (D). (D) Minimal mechanism for type 1 pilus rod assembly and assembly inhibition *in vitro*. The rate constants for binding/dissociation of FimCA and FimCI to/from FimDN and for uncatalyzed FimA assembly were kept fixed during fitting. Rate constants for FimCA had been measured previously (Nishiyama and Glockshuber, 2010). The domains of FimD are denoted: N-terminal (D_N_), plug (PD), transmembrane (TMD) and C-terminal domains (D_C1_, D_C2_).

### A comprehensive, quantitative model of pilus rod assembly and stochastic assembly termination by FimI

To obtain detailed and quantitative information on the assembly termination activity of FimCI, FimD-catalyzed FimA assembly kinetics were recorded under conditions where pilus rod assembly was initiated by addition of mixtures of FimCA and FimCI to activated FimDCH. Specifically, the initial FimDCH and FimCA concentrations were kept constant (0.1 μM and 20 μM, respectively) and the initial FimCI concentration was varied between 0.01 and 1 μM (Figure 2C). Indeed, FimA assembly was slowed down progressively with increasing FimCI concentrations. At the highest FimCI concentration tested (1 μM, 10-fold excess over FimD), FimCA assembly could no longer be detected during 65 minutes, analogous to incubation of FimCA alone (without FimDCH) (Figure S3A and S3B). The uncatalyzed FimCA polymerization proceeded extremely slowly and only yielded small FimA oligomers (but no rods) and was analyzed according to an irreversible second-order reaction as previously described (Vetsch et al., 2006), yielding the rate constant for spontaneous, uncatalyzed polymerization of FimCA k_(A-A)uncat_ = 0.73 ± 0.02 M^−1^ s^−1^.

The minimal kinetic model of FimD-catalyzed incorporation of an individual FimC-bound subunit into the growing pilus includes three rate constants: Binding (k_on_) and dissociation (k_off_) of the incoming FimC-subunit complex to the N-terminal periplasmic domain of FimD (FimD_N_) and irreversible subunit incorporation and displacement of the FimC molecule that initially capped the growing pilus end (k_DSE_). This is formally equivalent to a Michaelis-Menten-type model (Allen et al., 2013; Nishiyama et al., 2008). For global fitting of the kinetics of FimCA consumption at different FimCA/FimCI ratios (Figure 2C), we took into account that incorporation of the very first FimA subunit, i.e., binding to FimF at the pilus base, is slower than that of any subsequent FimA, where FimA-FimA contacts instead of a FimA-FimF contact are formed (Nishiyama et al., 2008). In addition, we included the small contribution of the very slow, FimD-independent (uncatalyzed) formation of FimA oligomers to FimCA consumption during the time window of our *in vitro* assembly experiment (65 min). Based on our systematic FimD activation experiments (Figure S2) and the previously described rate constants of FimD-catalyzed tip fibrillum assembly (Allen et al., 2013), we assumed that all activated FimD molecules carried a FimC-capped FimF at the growing end of the tip fibrillum. Regarding FimCI incorporation and assembly termination, we assumed that incorporation of a single FimI completely terminates pilus assembly, and that FimI can only bind to a terminal FimA subunit and not to FimF in the activated FimDHGFC complex (Figure 2D).

To minimize the number of variable parameters in the global fitting of the kinetics of FimCA consumption (Figure 2C), we determined the still unknown rate constants of association (k_on_) and dissociation (k_off_) for FimCI binding to FimD_N_ using stopped-flow fluorescence, as described previously (Nishiyama and Glockshuber, 2010) (Figure S3C). Similar to the binding of all other chaperone-subunit complexes to FimD_N_ (Nishiyama and Glockshuber, 2010), binding of FimCI to FimD_N_ proved to be very rapid (k_on_ = 1.45 · 10^8^ M^−1^ s^−1^), close to the diffusion-limit of ca. 10^9^−10^10^ M^−1^ s^−1^ (Berg and von Hippel, 1985; v. Smoluchowski, 1917), and highly dynamic (k_off_ = 94 s^−1^). Notably, the results showed that FimCI bound faster to and dissociated slower from FimD_N_ than any other FimC-subunit complex (Table 1).

**Table 1.** Kinetic constants and apparent specificity constants for FimD-catalyzed assembly of FimC-subunit complexes at pH 8.0.

| FimD substrate | $k_{on}$ ($M^{-1} s^{-1}$)<sup>a</sup> | $k_{off}$ ($s^{-1}$)<sup>a</sup> | $k_{DSE}$ ($s^{-1}$)<sup>b</sup> | $K_M^{app} (= \frac{k_{off} + k_{DSE}}{k_{on}})$ ($\mu M$) | $k_{DSE}/K_M^{app}$ ($M^{-1} s^{-1}$) | $\frac{k_{DSE}/K_M^{app}}{k_{DSE}/K_M^{app} (FimCl)}$ |
| --- | --- | --- | --- | --- | --- | --- |
| FimCl | $1.45 \cdot 10^8$ | 94 | 0.14 (I-A) | 0.65 | $2.2 \cdot 10^5$ | 1.0 |
| FimCG | $2.85 \cdot 10^7$ | 855 | 2.85 (G-H) | 30.1 | $9.5 \cdot 10^4$ | 0.44 |
| FimCA | $1.1 \cdot 10^7$ | 267 | 1.4 (A-A) | 24.4 | $5.7 \cdot 10^4$ | 0.27 |
| FimCF | $1.8 \cdot 10^7$ | 142 | 0.053 (F-G) | 7.9 | $6.7 \cdot 10^3$ | $3.1 \cdot 10^{-2}$ |
| FimCA | $1.1 \cdot 10^7$ | 267 | 0.0055 (A-F) | 24.3 | $2.3 \cdot 10^2$ | $1.1 \cdot 10^{-3}$ |

<sup>a</sup> $k_{on}$ and $k_{off}$ values refer to binding of FimC-subunit complexes to FimD<sub>N</sub>. Values for FimCG were taken from (Allen et al., 2013), values for FimCA and FimCF from (Nishiyama et al., 2008).
<sup>b</sup> $k_{DSE}$ values refer to formation of preferred subunit-subunit contacts, i.e. incorporation of FimG into FimH (G-H), FimF into FimG (F-G), FimA into FimF (A-F), FimA into FimA (A-A) and FimI into FimA (I-A).

With the experimentally determined k_on_ and k_off_ values for binding of FimCA and FimCI to FimDN and the rate constant for uncatalyzed polymerization of FimCA (k_(A-A)uncat_) as fixed parameters, all FimA assembly kinetics (Figure 2C) were then fitted globally according to the model in Figure 2D. As FimDCH preparations have been shown to contain a small fraction of inactive molecules that are unable to assemble a tip fibrillum (Allen et al., 2013), the total concentration of catalytically active FimDCH molecules was also included as an open fitting parameter. We obtained a value of 82 ± 1% for the fraction of active FimDCH molecules, in good agreement with the value of 90% reported earlier (Allen et al., 2013). As the global fit (solid lines in Figure 2C) did not show systematic deviations from the experimental data, we could complete our kinetic model of FimC-catalyzed pilus rod assembly (Figure 2D) with the DSE rate constants for irreversible binding of FimA to FimF (k_(A-F)_ = (5.5 ± 0.5)ˑ10^−3^ s^−1^), FimA to FimA (k_(A-A)_ = 1.4 ± 0.1 s^−1^) and FimI to FimA (k_(I-A)_ = 0.14 ± 0.01 s^−1^) (Table 1). The results confirmed that formation of a FimA-FimA contact is two orders of magnitude faster than binding of the first FimA to FimF (Nishiyama et al., 2008).

The FimC-subunit complexes FimCG, FimCF, FimCA and FimCI can be considered alternative substrates of FimD. Using the k_on_ and k_off_ values for binding of the different FimC-subunit complexes to FimD_N_ and the DSE rate constants k_(G-H)_, k_(F-G)_, k_(A-F)_, k_(A-A)_ and k_(I-A)_ determined in this study and previously (Allen et al., 2013; Nishiyama et al., 2008), we calculated their apparent K_M_ values 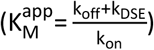 and specificity constants 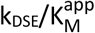 for the catalyst FimD (Table 1). Notably, the data showed that FimCI is the best substrate of all FimC-subunit complexes, even exhibiting a 4-fold higher specificity constant than FimCA. Overall, the specificity constants obtained for the different FimD substrates (2.3ˑ10^2^ to 2.2ˑ10^5^ M^−1^ s^1^) are in the range of enzymes with medium catalytic proficiency (Table 1).

Negative-stain electron microscopy of samples taken from the assembly reactions (Figure 2C) after 65 minutes of incubation confirmed that, as expected for pilus rod assembly termination by stochastic incorporation of FimI, the length of the assembled pilus rods decreased with increasing FimCI concentrations when FimCA was kept constant (Figure 3A). Pilus length distributions revealed that the majority of the population gradually shifted to shorter lengths and consequently the median pilus length steadily decreased, from 3.8 · 10^2^ nm for catalyzed assembly in the absence of FimI to 0.4 · 10^2^ nm for assembly in presence of a 3.3-fold excess of FimCI over FimD (Figure 3B). Figures 3A and 3B also show that two different pilus rod populations were obtained when substoichiometric concentrations of FimCI relative to FimD were used (in particular at FimCI:FimD ratios of 0.4:1, 0.6:1 and 0.8:1), so that some pilus rods assembled via FimD could not be terminated by FimCI: Besides FimCI-terminated pili shorter than 200 nm, a second population of much longer pili with lengths of up to ^~^1700 nm was observed under these conditions. For a comparison, the length of pili assembled in the absence of FimCI did not exceed 700 nm (Figure 3B). We interpret this result such that a larger pool of FimCA complexes was available for pilus rod assembly via the remaining, active FimD molecules when all FimCI complexes had been consumed and had inhibited the other FimD molecules at the early stage of the assembly reaction. In fact, even at the lowest FimCI concentrations used, longer pili already began to appear. The fraction of pili with lengths above 600 nm increased from about 1.5% in absence of FimI to 6.5% and 18% at FimCI:FimD ratios of 0.1:1 and 0.2:1, respectively (Figure 3B).

**Figure 3.**
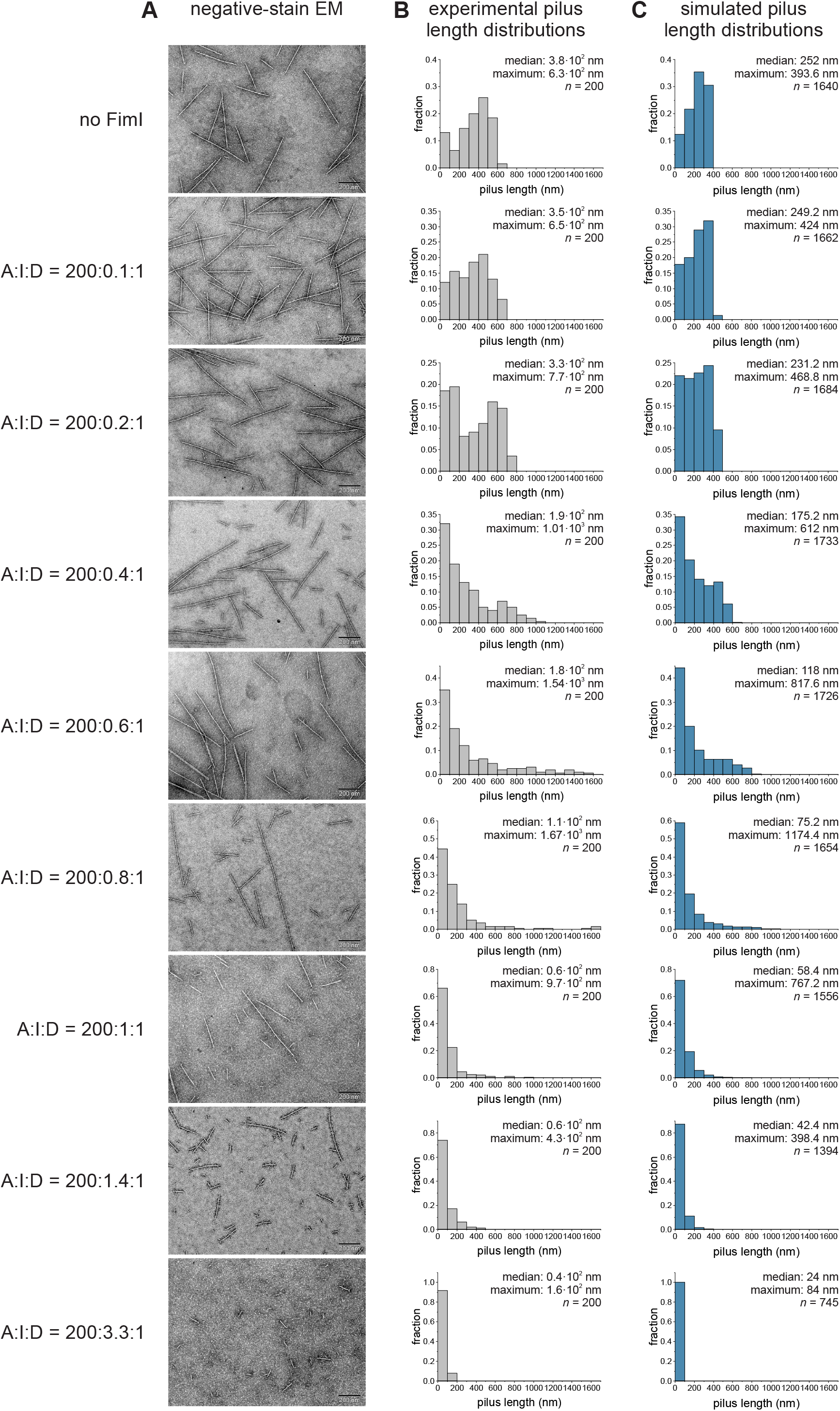
FimI modulates pilus length: Experimental *in vitro* data versus simulations *in silico*. (A) Representative negative-stain electron micrographs of samples taken from the pilus rod assembly reactions (Figure 2C) after 65 minutes of incubation (scale bar = 200 nm). The ratios between FimCA, FimCI and FimD (A:I:D) are indicated on the left and refer to the total initial concentrations of the respective proteins, not taking into account that only 82% of the FimD molecules were catalytically active. (B and C) Length distributions of pili assembled *in vitro* (grey) and *in silico* (blue). Median, maximum and size *n* of each sample are indicated.

Having a complete kinetic model of pilus rod assembly and assembly inhibition at hand (Figure 2D) allowed us to perform Monte Carlo simulations of FimD-catalyzed pilus rod assembly reactions at different FimCI/FimCA ratios and to generate simulated pilus length distributions for comparison with the experimental data (Figure 3C). The simulations were performed for the assembly reactions in Figure 2C, each with 2000 activated FimDHGFC molecules. Overall, pilus rods assembled *in silico* were found to be slightly shorter compared to the experimental data (median length values were between 57% and 93% of those found experimentally). Nevertheless, the simulated length distributions were qualitatively in full agreement with the experimental distributions in that the median length consistently decreased with increasing FimCI concentration (from 252 nm for assembly in the absence of FimCI to 24 nm for assembly in presence of a 3.3-fold excess of FimCI over FimD). In addition, at substoichiometric concentrations of FimCI relative to FimD, a fraction of up to 19% of the pili exceeded the maximum length of ^~^400 nm in the simulated length distribution in absence of FimCI (Figure 3B,C; see also above). Thus, within experimental error, the Monte Carlo predictions are in full agreement with our mechanistic model of pilus rod assembly termination by stochastic incorporation of FimI.

Furthermore, the simulation of pilus rod assembly made it possible to distinguish FimI-terminated pilus rods from non-terminated rods. The simulated length distributions, which included all pili, could therefore be deconvoluted into their two underlying subdistributions of pili with or without FimI bound (Figure S4). As one would expect, at the lowest FimCI:FimD ratio of 0.12:1, the entire distribution was still dominated by pilus rods that had no FimI bound. At higher FimCI:FimD ratios of 0.49:1 and 0.73:1, however, the vast majority of the pili shorter than 200 nm had FimI bound (80% and 95%, respectively), while most pili longer than 200 nm still had no FimI bound (90% and 72%, respectively). This observation is in agreement with FimCI being the best FimD substrate (Table 1), as this favors formation of predominantly short pili terminated with FimI (note that the fraction of pili shorter than 100 nm is always highest in all the subdistributions of FimI-terminated pili (Figure S4)). The simulations also confirmed that those rods that were both i) assembled at substoichiometric amounts of FimCI relative to FimD and ii) exceeded the maximum rod length of ^~^400 nm for assembly simulated in absence of FimI (see topmost panel in Figure 3C), were predominantly not capped by the assembly terminator FimI. Importantly, these longer pili disappeared at FimCI:FimD ratios above 1:1 in both the experimental data and the simulations (Figure 3B and 3C), indicating that all active FimD molecules became inhibited in the presence of equimolar amounts of FimCI. The deconvoluted length distributions further supported this indication, showing that practically all pili were indeed capped by FimI at the FimCI:FimD ratios 1.2:1, 1.7:1 and 4.0:1 (Figure S4).

In summary, our data show that the natural pilus length distribution results from a stochastic chain termination reaction in which the ratio between FimCA complexes and the FimCI termination complex dictates the pilus length distribution, in addition to the molar FimCA:FimD and FimCI:FimD ratios. The mechanism of PapH, the subunit terminating assembly in the related P pilus system (Baga et al., 1987; Verger et al., 2006), is likely identical to that of FimI.

### *In vivo* modulation of pilus length via the FimCA:FimCI and FimCI:FimD ratios

Type 1 pili attached to the outer *E. coli* membrane show an average length of a few hundred nanometers [(Hahn et al., 2002; Mulvey et al., 1998) and below]. Our pilus length distribution analysis revealed that 91% of the pili remained shorter than 200 nm at a 140-fold excess of FimCA over FimCI (Figure 3B, second-last panel). Thus, despite the fact that transcription of the *fimI* and *fimA* genes is controlled by the same promoter (Schwan, 2011) (Figure S5A), for pilus assembly to proceed to the lengths observed *in vivo*, the concentration of FimCI in the periplasm must be at least two orders of magnitude lower than that of FimCA. Expression of the *fimI* and *fimA* genes is therefore expected to be differentially regulated.

A prerequisite for FimC-catalyzed subunit folding is the DsbA-catalyzed formation of a single, conserved disulfide bond in the structure of all subunits (Crespo et al., 2012) (the disulfide oxidoreductase DsbA is the only known catalyst of disulfide bond formation in the *E. coli* periplasm). Therefore, differences between the kinetics or yields of oxidative folding of FimI and FimA could offer one explanation for different periplasmic concentrations of FimCI and FimCA. We tested this hypothesis and determined rate constants for i) DsbA-catalyzed oxidation of reduced, unfolded FimI or FimA 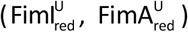 and ii) FimC-catalyzed folding of oxidized, unfolded FimI or FimA 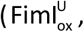 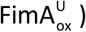 using stopped-flow tryptophan fluorescence spectroscopy experiments, essentially as described previously (Crespo et al., 2012) (Figure S5B-G). The kinetics of DsbA-catalyzed FimI oxidation measured under pseudo-first order conditions (6.7-fold excess of oxidized DsbA over 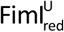) or using equimolar initial concentrations of oxidized DsbA and 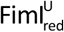 were fitted globally according to an irreversible, second-order reaction and yielded a rate constant of oxidation of 1.3 · 10^6^ M^−1^ s^−1^ (Figure S5B). In comparison, DsbA-catalyzed oxidation of 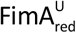 was 40-fold slower (3.4 · 10^4^ M^−1^ s^−1^, Figure S5E). FimC-catalyzed folding of 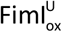 and 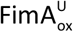 occurred with similar rates of 7.3 · 10^4^ M^−1^ s^−1^ and 4.3 · 10^4^ M^−1^ s^−1^ (Figure S5C and S5F). Consequently, the determined rate constants predicted similar *in vivo* half-lifes of ^~^0.4 and ^~^1 s for the formation of the native FimCI and FimCA complexes, respectively (Figure S5D and S5G). In agreement with previous results obtained for FimA, FimG and the pilin domain of FimH (FimH_P_) (Crespo et al., 2012), FimC-catalyzed folding of FimI was specific to disulfide-intact, unfolded FimI, as no fluorescence change was detected when FimC was mixed with 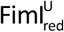 (Figure S5C). Moreover, DsbA- and FimC-catalyzed oxidative folding of both FimA and FimI proceeded to completion *in vitro*, without yield losses due to unspecific protein aggregation. Together, these results show that the kinetics of oxidative folding of FimI and FimA are very similar and therefore are unlikely to be responsible for differences in periplasmic FimCI and FimCA concentrations. To test whether different *fimI* and *fimA* transcript levels could account for the predicted ^~^10^2^-fold excess of FimCA over FimCI in the periplasm, we determined the relative *fimI* and *fimA* mRNA levels in *E. coli* W3110 cells by real-time PCR (Figure S5H). The *fimA* transcript level was only ^~^8-fold higher than that of *fimI* and ^~^12-fold higher than that of *fimC*, in good agreement with a previous study that used a DNA microarray to determine transcript levels in *E. coli* MG1655 cells (Schembri et al., 2002). We conclude that additional mechanisms likely contribute to the regulation of the periplasmic FimCA and FimCI concentrations, for example different efficiency of translation or co-translational translocation into the periplasm and/or proteolytic degradation.

The ≥100-fold higher periplasmic level of FimA relative to FimI implied by our *in vitro* assembly study is strikingly similar to a previous estimate for the PapA:PapH ratio (of at least 100:1) in P pilus biogenesis (Baga et al., 1987), supporting the idea that the general principles of pilus rod assembly termination and pilus length modulation are very similar for the type 1 and P pilus system. As a single type 1 pilus, on average, contains approximately 300 FimA molecules (see below and Table S1) and taking into account the predicted, more than 100-fold lower level of FimI, we can estimate the upper boundary of the periplasmic FimCI:FimD ratio to be roughly unity. Because DSE, and hence assembly inhibition, is irreversible, there is in fact no need for a much larger intracellular FimCI:FimD ratio to ensure quantitative assembly termination *in vivo*. It therefore appears that the regulation of the intracellular levels of FimD, FimCA and FimCI and the mechanism of FimD-catalyzed pilus biogenesis co-evolved with the natural type 1 pilus length distribution in *E. coli* (see Figure 4C).

**Figure 4.**
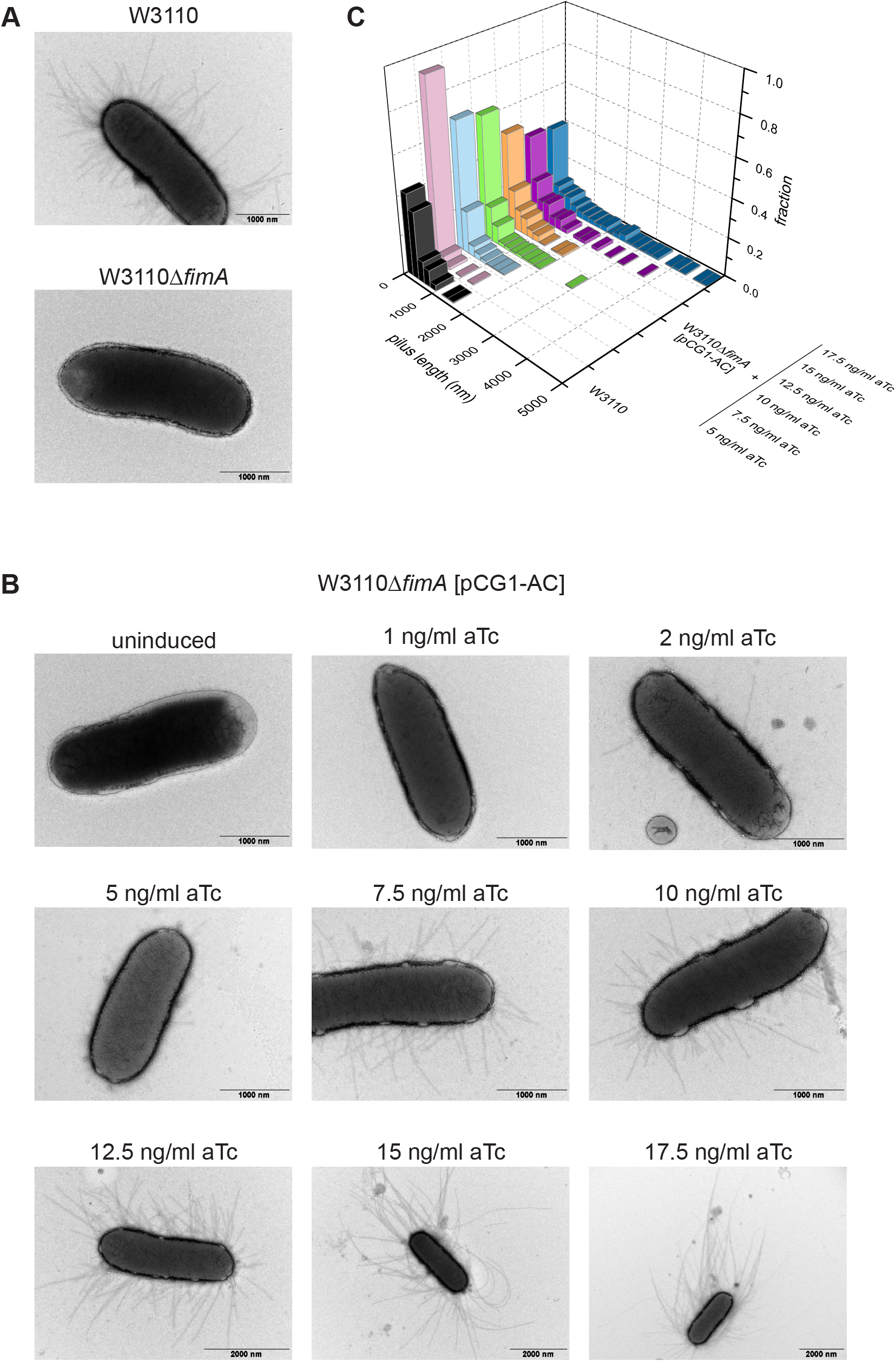
Modulation of type 1 pilus length *in vivo*. (A) Negative-stain electron micrographs of *E. coli* W3110 and W3110Δ*fimA*. (B) Negative-stain electron micrographs of *E. coli* W3110Δ*fimA* harboring plasmid pCG1-AC and grown in absence (uninduced) or presence of the indicated anhydrotetracycline (aTc) concentrations. (C) Pilus length distributions of W3110 and W3110Δ*fimA* [pCG1-AC] grown in presence of the indicated aTc concentrations (*n* = 200).

Next, we addressed the question of whether type 1 pilus length would directly depend on the FimCA:FimCI and FimCA:FimD ratios *in vivo*. Our above results predicted that the average length of pili displayed on cells would increase with increasing FimCA concentrations when the FimCI levels are kept constant. To test this hypothesis, we used the *fimA* deletion strain W3110Δ*fimA* (Hospenthal et al., 2017), in which base pairs 4‒528 of the *fimA* gene had been removed from the genome of *E. coli* W3110 wild-type. Negative-stain electron microscopy of W3110 and W3110Δ*fimA* cells confirmed that cells of the deletion strain no longer displayed type 1 pilus rods on their surface (Figure 4A). Successful deletion of *fimA* was also verified by real-time PCR (Figure S5H). To complement W3110Δ*fimA* cells and allow production of variable amounts of periplasmic FimCA, we constructed the plasmid pCG1-AC (Hospenthal et al., 2017) in which periplasmic coexpression of FimA with FimC is under control of the anhydrotetracycline-inducible *tetA* promoter, allowing fine-tuning of FimCA production by varying the concentration of the inducer anhydrotetracycline (aTc) in the growth medium (Neuenschwander et al., 2007).

W3110Δ*fimA* cells transformed with pCG1-AC and grown in presence of different aTc concentrations were then analyzed by negative-stain electron microscopy for their pilus length distributions. Figure 4B qualitatively shows that pilus length indeed increased with increasing aTc concentration. For a quantitative analysis, pili were released from cells by a heat step and length distributions were established for 200 pili in each preparation (Figure 4C and S6). While the median was 2.4 · 10^2^ nm for wild-type cells (corresponding to an average of ^~^300 copies of FimA per pilus), it increased from 0.7 · 10^2^ nm to 4.0 · 10^2^ nm for W3110Δ*fimA* [pCG1-AC] grown at aTc concentrations of 5 and 17.5 ng/ml aTc, respectively (Table S1). Similarly, the maximum pilus length (defined as the average length of the 5% longest pili) gradually increased from ^~^0.5 μm (5 ng/ml aTc) to ^~^4.0 μm (17.5 ng/ml aTc) (Table S1). These results show that pilus length in W3110Δ*fimA* [pCG1-AC] can be successfully modulated via the periplasmic FimCA levels (within a relatively narrow range of inducer concentrations). The FimCA:FimCI and FimCA:FimD ratios thus determine pilus length not only *in vitro* but also *in vivo*. Our results agree with a previous study on the related P pilus system, where periplasmic overproduction of the main subunit PapA or the terminating subunit PapH caused an increase or decrease in P pilus length, respectively (Baga et al., 1987). Similarly, while disruption of *mrpB*, encoding the terminating subunit MrpB of MR/P pili of *P. mirabilis*, led to significantly longer pili, overproduction of MrpB decreased pilus length in comparison to the parental wild-type strain (Li and Mobley, 1998).

### FimI contributes to stable anchoring of type 1 pili to the cell

In the P pilus system, the assembly terminator PapH was also shown to anchor P pili to the outer bacterial membrane (Baga et al., 1987). In light of the functional similarity between PapH and FimI suggested earlier (Rossolini et al., 1993) and established in this study, we hypothesized that FimI, like PapH, could be involved in anchoring type 1 pili to the bacterial outer membrane. To test this, an *E. coli* W3110 *fimI* deletion strain (W3110Δ*fimI*) was generated. Real-time PCR confirmed successful deletion of *fimI* (Figure S5H). W3110Δ*fimI* cells were then grown in either static or shaken liquid medium and probed for the presence of functional pili on the bacterial surface by testing the ability of W3110Δ*fimI* to agglutinate with yeast cells bearing highly mannosylated mannoproteins in the cell wall (Figure 5A). While agglutination readily occurred when W3110Δ*fimI* cells had been grown statically, no agglutination was detected with W3110Δ*fimI* grown in shaken medium, indicating that pili in W3110Δ*fimI* were sheared off mechanically under shaking and lost to the culture medium. A control showed that plasmid-encoded FimI complemented *fimI* deficiency of W3110Δ*fimI* and restored membrane anchoring and yeast agglutination.

**Figure 5.**
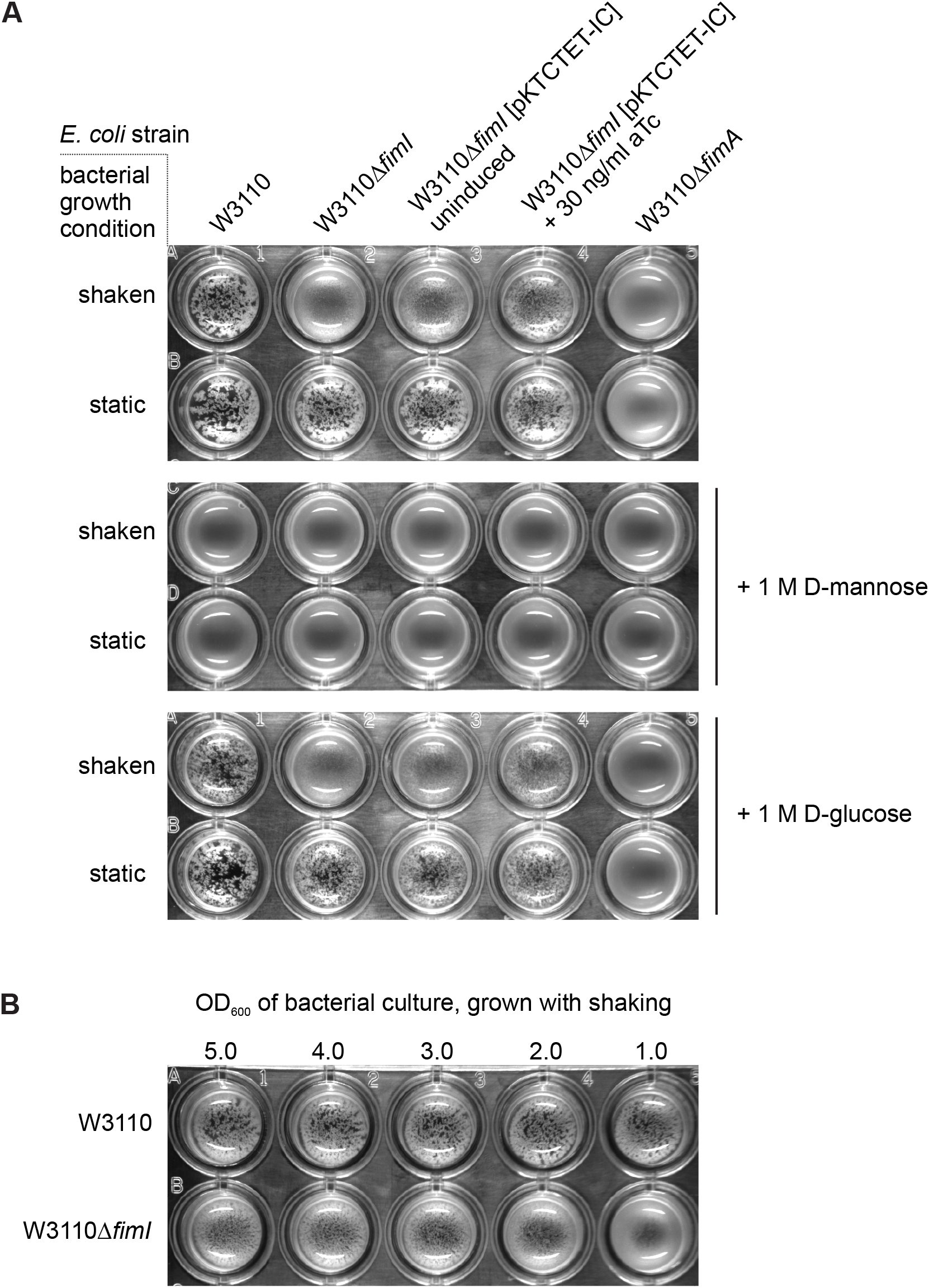
FimI anchors type 1 pili to *E. coli* cells. (A) Yeast agglutination assays performed with *E. coli* W3110, W3110Δ*fimI* and W3110Δ*fimA*, which had been grown under shaking or static conditions. To complement the *fimI* deletion, the plasmid pKTCTET-IC was used and gene expression induced with 30 ng/ml anhydrotetracycline (aTc). Mannose-specificity of yeast cell binding was tested by a competition assay in the presence of 1 M D-mannose or 1 M D-glucose (negative control). (B) Yeast agglutination assay with *E. coli* W3110 and W3110Δ*fimI*, grown under shaking conditions and diluted to the indicated OD_600_ values prior to mixing with the yeast cells.

As expected, wild-type W3110 cells displayed similar piliation levels for both growth conditions and caused agglutination even when grown under shaking. Notably, W3110Δ*fimI* cells grown under shaking also agglutinated weakly with yeast cells when used at higher cell densities in the assay (Figure 5B). This suggests that not all pili of W3110Δ*fimI* were lost to the culture medium during shaking, but that the loss of pili lowered the agglutination titer of the W3110Δ*fimI* strain compared to that of W3110. A control showed that addition of 1 M D-mannose blocked agglutination throughout, while addition of 1 M D-glucose (the C2 epimer of mannose) did not affect agglutination (Figure 5A). This confirms that the agglutination assay indeed reported on the presence of functional FimH at the tip of membrane-anchored type 1 pili. W3110Δ*fimA* cells failed to agglutinate yeast irrespective of whether the cells were grown statically or under shaking (Figure 5A). This agrees with a previous study that demonstrated a strongly impaired adhesion phenotype for *E. coli* cells with inactive *fimA* gene (Klemm et al., 1990). In the absence of FimA, tip fibrillae may thus still be assembled, but may be too short for reaching enough target mannosides on yeast cell walls. Together, the results show that FimI contributes to anchoring type 1 pili stably to the bacterial outer membrane under shear stress.

### Binding of FimI to the terminal FimA subunit slows dissociation of FimC from FimI 220-fold

The periplasmic, growing ends of type 1 pili are capped with a FimC chaperone that binds to the terminal pilus subunit (FimI in wild-type *E. coli* and FimA in Δ*fimI* strains) via donor strand complementation. The FimC (22.7 kDa) at the growing end is too large to pass the translocation pore of the assembly platform FimD. Release of a pilus to the extracellular medium would require dissociation of periplasmic FimC from the last incorporated pilus subunit. Therefore, FimI at the periplasmic pilus end could contribute to anchoring of the pilus to the outer membrane under mechanical stress by binding particularly strongly to FimC. Likewise, a significant fraction of membrane-associated pili may get lost to the surrounding medium in the absence of FimI. To address this question, we determined the rate constants of spontaneous dissociation of FimC_His_ (FimC with C-terminal hexahistidine tag) from either the binary FimC_His_-FimI (FimC_His_I) or the ternary FimC_His_-FimI-FimAt_His_ complex [(FimC_His_IA, (the “t” in FimAt_His_ indicates “N-terminal truncation, without donor strand”)] by measuring the time course of competitive displacement of FimC_His_ by excess untagged FimC (Vetsch et al., 2006). While FimC_His_I corresponds to the periplasmic preassembly state of FimI, the ternary FimC_His_IA complex represents the product of the terminal DSE reaction in type 1 pilus assembly.

As an homogeneous FimC_His_IA complex containing FimA wild-type was difficult to prepare due to FimA self-polymerization, we used a truncated FimA variant lacking the N-terminal donor strand (FimAt_His_) to obtain the homogeneous FimC_His_IA complex. FimC_His_I or FimC_His_IA (4 μM) were incubated in presence of a 9-fold molar excess of untagged FimC (36 μM) at 37 °C and the kinetics of attaining the new equilibrium were monitored with analytical cation exchange chromatography (Figure 6A and 6B). Notably, FimC_His_ dissociated from FimC_His_IA 220-fold slower than from FimC_His_I (with dissociation half-lifes of 3.85 ± 0.43 h and 63 ± 5.7 s for FimC_HisI_A and FimC_His_I, respectively), showing that binding of FimI to the last FimA subunit further stabilizes the FimI-FimC interaction. In addition, association between FimI and FimC_His_ is intrinsically tighter than that between FimA and FimC_His_, because FimC_His_ dissociates about 10-fold faster from FimA [half-life < 5 s (Vetsch et al., 2006)] than from FimI.

**Figure 6.**
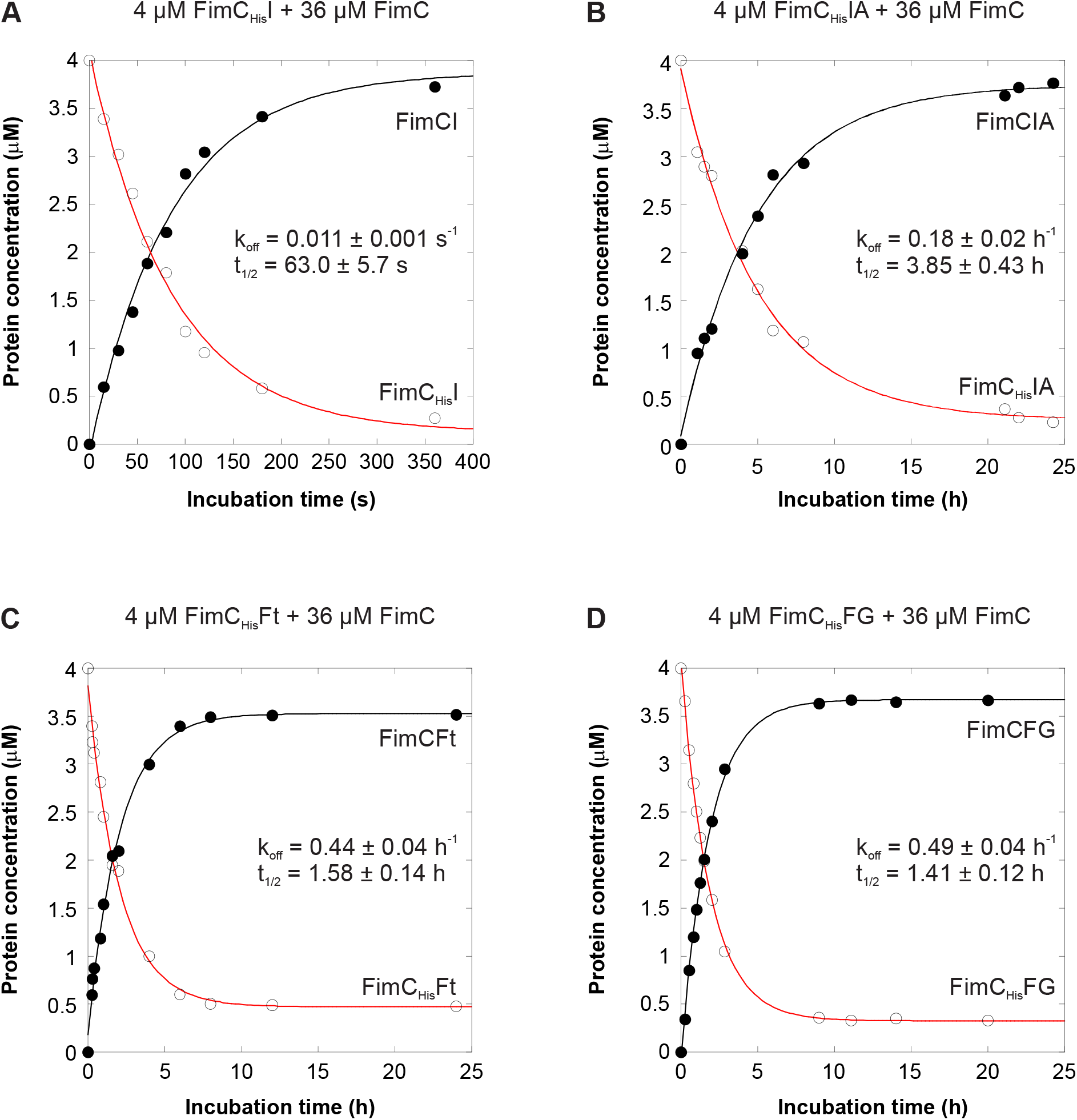
Kinetics of FimC_His_ dissociation from different chaperone-subunit complexes at 37 °C. For determination of the FimC_His_ dissociation rate constant (k_off_), 4 μM of FimC_His_I (A), FimC_His_IA (B), FimC_His_Ft (C) or FimC_His_FG (D) were incubated with a 9-fold molar excess of FimC. Relaxation to the new equilibrium was monitored by analytical cation exchange chromatography. Solid lines are fits according to a single-exponential function.

To test whether the slowed FimC_His_ dissociation upon FimI-FimA contact formation in FimC_His_IA was a unique property of FimC_His_IA or is generally observed when a binary chaperone-subunit complex associates with another subunit to a ternary chaperone-subunit-subunit complex, we also measured FimC_His_ dissociation rates for the FimC_His_-FimFt (FimC_His_Ft) and FimC_His_-FimF-FimGt (FimC_His_FG) complexes as a control (Figure 6C and 6D). For both complexes, we obtained practically identical FimC_His_ dissociation half-lifes (1.58 ± 0.14 h and 1.41 ± 0.12 h, respectively). Thus, in contrast to contact formation between FimI and FimA in FimC_His_IA, contact formation between FimF and FimG in FimC_His_FG does not slow FimC_His_ dissociation from FimG, possibly hinting at an allosteric stabilization of FimC_His_I against dissociation via FimI-FimA contact formation.

### Crystal structures of the FimC_His_-FimIt, FimD_N_-FimC_His_-FimIt and FimC_His_-FimI-FimAt_His_ complexes

As a step towards understanding the structural basis for termination of type 1 pilus assembly by FimI, we solved the crystal structures of the FimC_His_-FimIt (FimC_His_It), FimD_N_-FimC_His_-FimIt (FimD_N_CI) and FimC_His_-FimI-FimAt_His_ (FimC_His_IA) complexes to 1.75, 1.7 and 2.8 Å resolution, respectively (Figure 7A, 7B, 7D). In all three structures, the overall folds of FimC and FimI are very similar [Cα root-mean-square deviation (RMSD) values were 1.2 Å (superimposing FimC_His_It with FimCI of FimD_N_CI), 0.92 Å (superimposing FimC_His_It with FimC_His_I of FimC_His_IA) and 0.95 Å (superimposing FimC_His_I of FimC_His_IA with FimCI of FimD_N_CI)]. Similarly, the overall structure of FimD_N_CI closely resembled that of two other ternary FimD_N_-FimC-subunit complexes, FimD_N_CF and FimD_N_CH_P_ (Eidam et al., 2008; Nishiyama et al., 2005), with Cα RMSD values of 1.4 Å (superimposing FimD_N_CI and FimD_N_CH_P_), 1.4 Å (superimposing FimD_N_CF and FimD_N_CH_P_) and 1.2 Å (superimposing FimD_N_CI and FimD_N_CF) (Figure 7C). Given the high structural similarity of FimC and FimI in the FimC_His_It, FimD_N_CI and FimC_His_IA complexes, we focus on the analysis of the FimC_His_IA structure in the following.

**Figure 7.**
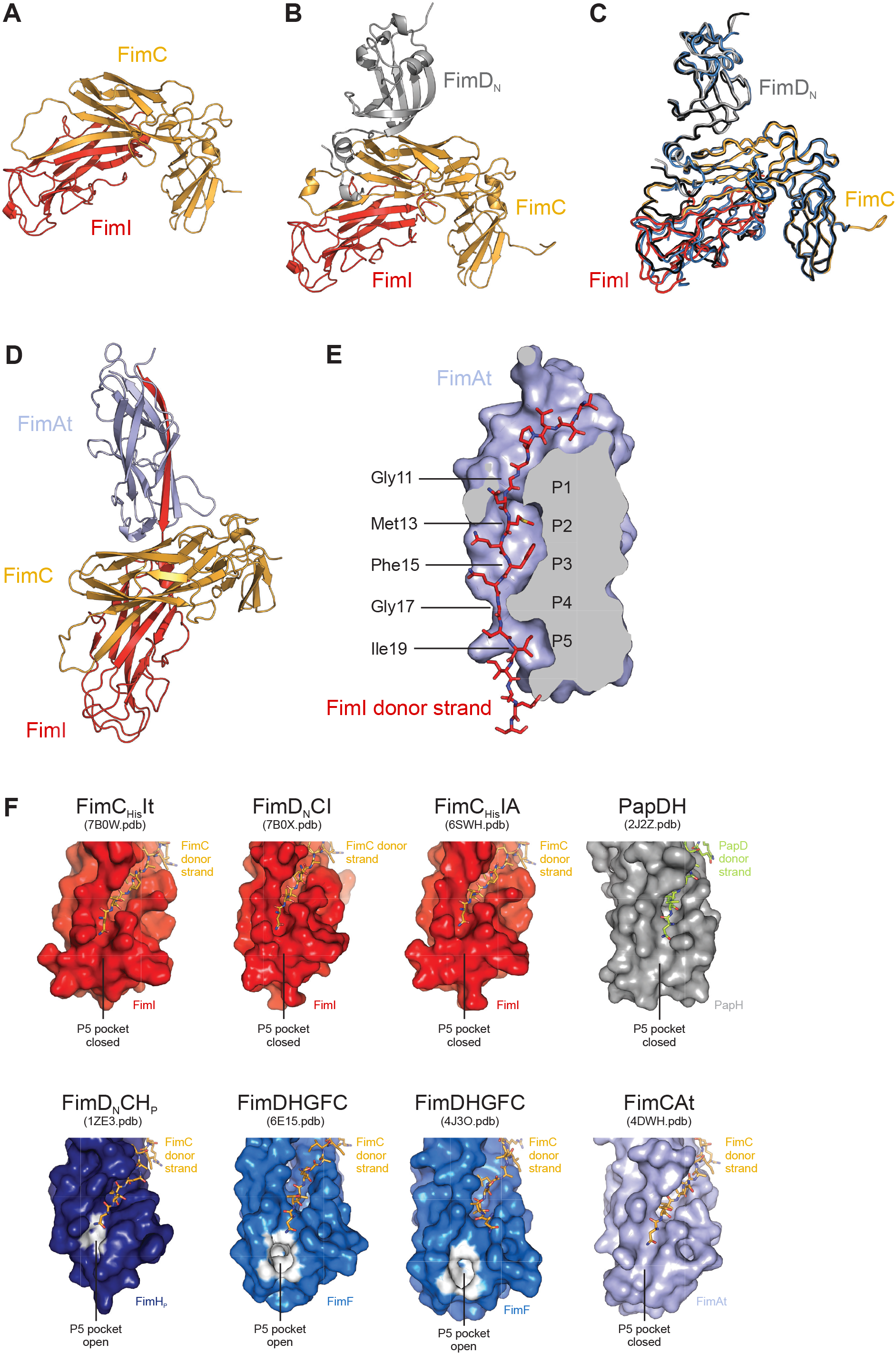
Crystal structures of the FimC_His_It, FimD_N_CI and FimC_His_IA complexes. (A) Crystal structure of the binary FimC_His_It complex. Proteins are shown in cartoon representation with FimC in gold and FimI in red. (B) Crystal structure of the ternary FimD_N_CI complex. Proteins are shown in cartoon representation with FimD_N_ in grey, FimC in gold and FimI in red. (C) Superimposition of the structures of the FimD_N_CI, FimD_N_CF and FimD_N_CH_P_ complexes. FimD_N_CI is colored as in panel (B), FimD_N_CF is in blue and FimD_N_CH_P_ in black. (D) Crystal structure of the ternary FimC_His_IA complex. All proteins are shown in cartoon representation with FimC in gold, FimI in red and FimAt in light blue. (E) Complementation of the FimAt fold by the FimI donor strand. FimAt is shown in sliced surface representation, the FimI donor strand is shown as stick model. P1 to P5 indicate the positions in the hydrophobic groove of FimAt where side chains of the FimI donor strand interact with the core of FimAt. (F) Conformation of the P5 pocket in FimI, PapH, FimH_P_, FimF or FimAt as observed in the structures of FimC_His_It, FimD_N_CI, FimC_His_IA, PapDH, FimD_N_CH_P_, the complex between FimD and the type 1 pilus tip (FimDHGFC), and FimCAt. While analysis of the structures by the CASTp web server (Tian et al., 2018) did not reveal a distinct P5 pocket in FimI, PapH and FimAt, the P5 pocket is clearly defined in FimHP of the FimD_N_CH_P_ complex and in FimF of the FimDHGFC complex. All subunits are shown in surface representation. FimH_P_ and FimF atoms identified by CASTp to be involved in formation of the P5 pocket are highlighted in white.

The asymmetric unit in the FimC_His_IA crystals contained two copies of the FimC_His_IA complex with very similar structures (RMSD of 1.4 Å for 458 aligned Cα atoms). As complex 1 (chains A, B and C) exhibited disorder near the N-terminus of FimAt and in some loop regions, complex 2 (chains D, E and F) was used for structural analysis. The interactions between FimC and FimI in the FimC_His_IA complex were found to be analogous to those observed in the complexes of FimCH, FimCF and FimCA (Crespo et al., 2012; Eidam et al., 2008; Nishiyama et al., 2005). Specifically, FimC completes the FimI fold by DSC and inserts residues 101‒110 of its G1-strand between the A’’- and F-strand of FimI in a parallel orientation with respect to the latter. Leu103, Leu105 and Ile107 of FimC interact with the hydrophobic core of FimI and an additional 22 intermolecular hydrogen bonds, 14 of which are hydrogen bonds between main-chain atoms, are formed between the FimC G1-strand and FimI. Furthermore, FimC residues 1‒ 7 interact with FimI residues 21-26 via six intermolecular hydrogen bonds, two of which are main-chain, and both Arg8 and Lys112 of FimC form a salt bridge with the C-terminus of FimI.

FimI itself possesses the characteristic, incomplete Ig-like fold with six β-strands, a short α-helical segment (residues 66‒69) and a secondary structure topology similar to that of other type 1 or P pilus subunits (Figure S8A and S8B) (Crespo et al., 2012; Eidam et al., 2008; Nishiyama et al., 2005; Puorger et al., 2008; Verger et al., 2007; Verger et al., 2006). Consequently, the Ig-like fold of FimI superimposes well onto that of the FimH pilin domain (FimH_P_), FimF, FimG, FimA and PapH, with pairwise Cα RMSD values between 1.5 and 1.8 Å (Figure S8C). The first resolved residue near the FimI N-terminus in the FimC_His_IA structure was Thr6. Residues 6‒20 of FimI constitute the FimI donor strand and insert into the hydrophobic groove between the A- and F-strand of FimAt in an antiparallel orientation compared to FimAt’s F-strand. Met13, Phe15 and Ile19 of FimI point towards the center of FimAt and complete its hydrophobic core by occupying binding pockets P2, P3 and P5, respectively (Figure 7E). Similar to what had been observed in structures of FimA complemented with the FimA donor strand (Puorger et al., 2011; Walczak et al., 2014), the FimA pockets P1 and P4 are shallow and hence occupied by Gly11 and Gly17 of FimI (Figure 7E). In addition to these interactions, 31 intermolecular hydrogen bonds are formed between the FimI donor strand and FimAt (24 of which are between main-chain atoms), and Glu22 of FimI interacts with Thr39 and Ala40 of FimAt through another four hydrogen bonds.

Besides these FimC-FimI and FimI-FimAt contacts, the FimC_His_IA complex harbors a third interface. It is located between FimC and FimAt, has an interface area of 351 Å^2^ and involves both domains of FimC that form contacts to the loops connecting β-strands A and B, C and D, and E and F of FimAt (Figure S8D). In this interface, FimC and FimAt interact via a salt bridge between Glu62 of FimC and Arg38 of FimAt, a single hydrogen bond between the side chain of Asn191 of FimC and the main-chain carbonyl oxygen of Ala93 of FimAt, and through hydrophobic interactions between Ala195 of FimC and Pro145 of FimAt (Figure S8D). These additional FimC-FimAt interactions may be the reason for the 220-fold slower dissociation of FimC_His_ from FimC_His_IA compared to FimC_His_I, and may prevent the dissociation of FimC from FimI at the pilus base and hence anchor the entire pilus to the membrane.

Previously, X-ray and electron cryomicroscopy (cryo-EM) structures of a pilus assembly intermediate, the complex between FimD and the type 1 pilus tip fibrillum (FimDHGFC), were solved (Du et al., 2018; Geibel et al., 2013). In these structures, FimH is already translocated to the extracellular side of the outer membrane, FimG is located inside the FimD pore, and FimF and FimC are bound to the CTDs on the periplasmic side of FimD. On this structural basis and together with the solved FimC_His_IA structure, we tried to model the FimD-bound structure of the proximal end of the type 1 pilus rod terminated by FimCI. Assuming that FimCI, after donor strand exchange, is transferred from FimD_N_ to the CTDs of FimD, we superimposed FimC_His_ of the FimC_His_IA complex onto FimC of the FimDHGFC complexes. While the structures of FimI and FimF superimposed rather well, FimAt of the FimC_His_IA complex did not line up with FimG, but clearly clashed with FimD’s transmembrane domain (Figure S8E). This discrepancy suggests that either FimD changes its conformation upon incorporation of FimA and formation of the helical quaternary structure of the rod on the extracellular side of FimD (the structure of FimD in complex with an assembled pilus rod is still unknown) or conformational changes occur within FimD and/or FimCIA upon assembly termination by incorporation of FimCI.

For the P pilus system, previous work suggested that assembly termination by PapH is caused by the absence of a P5 pocket in the PapDH structure (Verger et al., 2006), thus making PapH incapable of accepting donor strands of incoming chaperone-subunit complexes. Analysis of the FimC_His_IA crystal structure by the CASTp web server (Tian et al., 2018) revealed that FimI, akin to PapH, does not possess a distinct P5 pocket (Figure 7F). The same analysis revealed that the P5 pocket is in an open conformation in FimH_P_ of the FimD_N_CH_P_ complex (with a volume of 3 Å^3^), as well as in FimF in the structures of the FimDHGFC complex with volumes of 51.3 Å^3^ and 18.4 Å^3^ for the cryo-EM and crystal structure, respectively (Figure 7F). Therefore, both FimI and PapH may indeed achieve termination of pilus rod assembly by analogous mechanisms that prevent displacement of FimC at the growing pilus end by an incoming donor strand. Notably, however, the P5 pocket of FimA in the structure of the FimCAt complex is also closed (Figure 7F). Whether or not there is a clear-cut correlation between an open/closed P5 pocket in the FimC-capped subunit at the pilus base and its ability to undergo DSE with another subunit thus still remains to be shown.

In summary, we here identified FimI as the terminating subunit in type 1 pilus rod assembly and presented a quantitative model for rod assembly and its termination that is consistent with the natural pilus length distribution and in full agreement with a stochastic chain termination mechanism. Our results, in particular the *in vitro* reconstitution of FimD-catalyzed pilus rod assembly and termination, provide a basis for structure determination of FimD bearing an assembled pilus rod on the extracellular side and the FimCI termination complex on the periplasmic side by cryo-EM. The *in vitro* reconstitution of type 1 pilus assembly also contributes to a better understanding of enzyme-catalyzed assembly of filamentous protein polymers and provides a general framework for testing the mechanism of assembly termination and pilus length regulation in related filamentous pilus systems.

## ACKNOWLEDGMENTS

We thank the lab of Donald Hilvert and Peter Kast for generously providing plasmid pKTCTET-0; Gabriel Waksman for kindly providing plasmids pAN2-Strep and pETS1001; Marcel Bolten, Marc Leibundgut and Lena Keller for discussions; the Scientific Center for Optical and Electron Microscopy (ScopeM) of ETH Zurich for providing access to the electron microscope, and Peter Tittmann for technical support. We are grateful to Stephan Handschin from ScopeM for help with Fiji and TrakEM2; to Andrea Prota for help with refinement; to Arthur Goldsipe for help with stochastic simulations; to Jessica Stanisich for critical reading of the manuscript, and to Hiang Dreher-Teo and Helene Fäh-Rechsteiner for excellent technical assistance. This work was supported by grants 31003A_156304 and 310030B_176403/1 from the Swiss National Science Foundation to R.G.

## AUTHOR CONTRIBUTIONS

C.G. reconstituted pilus assembly, measured and evaluated kinetic data, performed electron microscopy, generated experimental and simulated pilus length distributions, performed *in vivo* titration of FimA and agglutination experiments, and designed experiments; C.P. performed and evaluated kinetic experiments; O.I. measured and evaluated binding of FimCI to FimD_N_, determined FimC_His_ dissociation rates and crystallized FimC_His_IA; Z.B. investigated oxidative folding of FimI and FimA, performed real-time PCR experiments and crystallized FimC_His_It and FimD_N_CI; M.E.W. measured kinetic data; M.A.S. and G.C. solved the structures of FimC_His_It, FimD_N_CI and FimC_His_IA; R.G. designed experiments and supervised the work; C.G. and R.G. wrote the paper, with contributions from C.P., O.I., Z.B. and M.A.S.

## METHODS

### Bacterial strains and plasmids

The *E. coli* W3110Δ*fimI* strain was created by deleting amino acids 43-116 of the mature FimI protein as described (Link et al., 1997) using wild-type *E. coli* W3110 cells (Bachmann, 1972) as parental strain and, unintentionally as a result of the cloning strategy, replacing this amino acid stretch by the sequence CLSLVDG.

The DNA sequence encoding FimI without its signal peptide was amplified by PCR using genomic DNA of *E. coli* W3110 cells as template and cloned into pET-11a (Studier et al., 1990) yielding the plasmid pFimI_cyt where *fimI* transcription is controlled by the T7*lac* promoter. For periplasmic coexpression of *fimI* and *fimC*, both genes were subcloned from plasmid pACIC-P_tet_ (Ignatov, 2009) into pKTCTET-0 (Roderer et al., 2014) via NdeI and SpeI restriction sites. The resulting plasmid was termed pKTCTET-IC and contains a *tetA/T7* tandem promoter, the *tetR*, *bla*, *fimI* and *fimC* genes and a pUC origin of replication. Plasmids for periplasmic expression of FimCG and FimCF were obtained by cloning the genes encoding FimG and FimC or FimF and FimC into pTrc99A (Amann et al., 1988). Plasmid pfimC_cyt for cytoplasmic expression of FimC was generated by subcloning the *fimC* gene from pfimC_His__cyt (Vetsch et al., 2004) into pET-11a via NdeI and BamHI restriction sites.

### Protein production

The N-terminal domain of FimD (FimD_N_, residues 1-125 of FimD), a FimC variant with a C-terminal hexahistidine tag (FimC_His_) and the FimC_His_-FimF (FimC_His_F), FimC_His_-FimFt (FimC_His_Ft) and FimC_His_-FimGt (FimC_His_Gt) complexes were expressed and purified as described (Gossert et al., 2008; Nishiyama et al., 2005; Nishiyama et al., 2003; Vetsch et al., 2004). FimAt_His_ (wild-type FimA missing residues 1-13 but with an N-terminal hexahistidine tag) was expressed and purified as described (Puorger et al., 2011).

The ternary complex between FimD, FimC and FimH (FimDCH) was produced as described (Geibel et al., 2013; Phan et al., 2011). Briefly, *E. coli* Tuner cells carrying pAN2-Strep and pETS1001 were grown at 37 °C in TB medium containing kanamycin at 30 μg/ml and spectinomycin at 100 μg/ml. At OD_600_ = 1.0, gene expression was induced by adding IPTG to 100 μM and L-arabinose to 0.1% (w/v). Glycerol was added to 0.1% (v/v) and the cells grown for 48 h at 16 °C. Cells were harvested by centrifugation, resuspended in 20mM Tris-HCl pH 8.0 (30 ml per liter of culture) containing complete EDTA-free protease inhibitor cocktail (Roche) and lysed using a Microfluidizer M-110L (Microfluidics, USA). After centrifugation (10 min, 5000xg, 4 °C), the supernatant was recovered, sarkosyl added to 0.5% (w/v) and the solution stirred for 5 min at room temperature. Outer membranes were pelleted by ultracentrifugation (1 h, 100000xg, 4 °C) and resuspended with 9 ml of 20 mM Tris-HCl pH 8.5, 120 mM NaCl supplemented with protease inhibitors (Roche) per gram of membranes using a dounce homogenizer. For solubilization, n-dodecyl-β-D-maltopyranoside (DDM) was added to 1.5% (w/v), the suspension stirred for 30 min at room temperature and insoluble material removed by ultracentrifugation (45 min, 100000xg, 4 °C). The supernatant was passed over a 5 ml HisTrap HP column (GE Healthcare) equilibrated with 20 mM Tris-HCl pH 8.5, 120 mM NaCl, 0.05% (w/v) DDM (buffer A). The column was washed with buffer A containing 25 mM imidazole and bound protein eluted by a step gradient with buffer A containing 250 mM imidazole. The solution was diluted 2-fold with buffer A, loaded onto an 8 ml Strep-Tactin sepharose column (IBA GmbH) equilibrated with buffer A, washed with buffer A and bound protein eluted with buffer A containing 2.5 mM D-desthiobiotin. The solution was concentrated by ultrafiltration using Amicon Ultra centrifugal filters with 100 kDa molecular weight cutoff (Merck) and passed over a Superdex 200 26/60 gel filtration column (GE Healthcare) equilibrated with 20 mM Tris-HCl (pH 8.0 at room temperature), 50 mM NaCl and 0.05% (w/v) DDM. FimDCH eluted in two main peaks, of which the last one was pooled, concentrated to approx. 1 μM, aliquoted, flash-frozen in liquid nitrogen and stored at −80 °C.

For the production of the FimC-FimF (FimCF) and FimC-FimG (FimCG) complexes, *E. coli* HM125 cells carrying pfimF-C-ATG-trc or pfimGC-trc were grown at 30 °C in 2YT medium containing ampicillin at 100 μg/ml. At OD_600_ = 0.7, expression was induced by adding IPTG to 1 mM and the cells grown further for 4 h. Cells were harvested by centrifugation (10 min, 4200xg, 4 °C), resuspended in 50 mM Tris-HCl pH 7.5, 150 mM NaCl, 5 mM EDTA, 1 mg/ml polymyxin B sulfate (10 ml buffer per liter of culture) and shaken for 1 h at 4 °C. After centrifugation (30 min, 48000xg, 4 °C), the supernatant was dialyzed against 20 mM Tris-HCl pH 8.0, centrifuged (15 min, 48000xg, 4 °C) and loaded onto a 15 ml Q-sepharose FF (GE Healthcare) column equilibrated with the same buffer. For FimCF, the flowthrough was collected and solid ammonium sulfate added to a final concentration of 1.2 M. After centrifugation (10 min, 48000xg, 4 °C), the solution was loaded onto a 10 ml Phenyl-sepharose HP (GE Healthcare) column equilibrated with 20 mM Tris-HCl pH 8.0, 1.2 M (NH_4_)_2_SO_4_ and bound protein eluted with a linear gradient from 1.2 – 0 M (NH_4_)_2_SO_4_. Fractions containing FimCF were pooled, dialyzed against 20 mM MES-NaOH pH 5.5, loaded onto a Source 30S (GE Healthcare) column equilibrated with the same buffer and proteins eluted with a linear gradient from 0-200 mM NaCl. FimCF typically eluted at a conductivity of approx. 8 mS/cm. The appropriate fractions were pooled, the solution concentrated by ultrafiltration using Amicon Ultra centrifugal filters with a 10 kDa molecular weight cutoff (MWCO) (Merck) and passed over a Superdex 75 16/60 gel filtration column (GE Healthcare) equilibrated with 20 mM Tris-HCl (pH 8.0 at room temperature), 50 mM NaCl.

Fractions containing pure FimCF complex were pooled, aliquoted, flash-frozen in liquid nitrogen and stored at −80 °C. FimCG was eluted from the Q-sepharose FF column with a linear gradient from 0-400 mM NaCl. Fractions containing the majority of FimCG were pooled, dialyzed against 20 mM MOPS-NaOH pH 6.7 and loaded onto a 6 ml Resource S column (GE Healthcare) equilibrated with the same buffer. Bound FimCG was eluted with a linear gradient from 0-200 mM NaCl, fractions containing FimCG were pooled and solid ammonium sulfate added to a final concentration of 1.2 M. After centrifugation (10 min, 48000xg, 4 °C), the solution was loaded onto a 10 ml Phenyl-sepharose HP (GE Healthcare) column equilibrated with 20 mM MOPS-NaOH pH 6.7, 1.2 M (NH_4_)_2_SO_4_ and bound protein eluted with a linear gradient from 1.2 – 0 M (NH_4_)_2_SO_4_. Fractions containing FimCG were pooled, the solution concentrated by ultrafiltration using Amicon Ultra centrifugal filters with 10 kDa molecular weight cutoff (Merck) and passed over a Superdex 75 26/60 gel filtration column (GE Healthcare) equilibrated with 20 mM Tris-HCl (pH 8.0 at room temperature), 50 mM NaCl. Fractions containing pure FimCG complex were pooled, aliquoted, flash-frozen in liquid nitrogen and stored at −80 °C. Final yields were 0.5 and 1.7 mg per liter of culture for FimCF and FimCG, respectively. The identity of the proteins was confirmed by ESI-MS: Expected/measured masses for FimC were 22730.1/22729.5 and 22729.0 Da; for FimF: 16166.2/16165.0 Da; and for FimG: 14854.3/14853.0 Da.

FimC was produced by growing *E. coli* BL21(DE3) cells harboring pfimC_cyt at 37 °C in 2YT medium containing ampicillin at 100 μg/ml. At OD_600_ = 1.0, expression was induced by adding IPTG to a final concentration of 1 mM and the cells grown further for 4 h. Cells were harvested by centrifugation (10 min, 4200xg, 4 °C), resuspended in 100 mM Tris-HCl pH 8.0, 1 mM EDTA (3 ml buffer per gram of cells) and lysed using a Microfluidizer. After centrifugation (45 min, 48000xg, 4 °C), the supernatant was dialyzed against 10 mM Tris-HCl pH 8.0, centrifuged (15 min, 48000xg, 4 °C) and passed over a Q-sepharose FF (GE Healthcare) column. The flowthrough was collected, dialyzed against 20 mM MES-NaOH pH 6.0, centrifuged as above and loaded onto a Source 30S (GE Healthcare) column equilibrated with the same buffer. Bound FimC was eluted with a linear gradient from 0-200 mM NaCl. Fractions containing FimC were pooled and solid ammonium sulfate added to a final concentration of 1.4 M. The protein solution was applied to a Phenyl-sepharose HP (GE Healthcare) column equilibrated with 20 mM MES-NaOH pH 6.0, 1.4 M (NH_4_)_2_SO_4_ and FimC eluted with a linear gradient from 1.4 – 0 M (NH_4_)_2_SO_4_. FimC-containing fractions were pooled, dialyzed against water, flash-frozen in liquid nitrogen and stored at −20 °C. The final yield was 40 mg per liter of culture.

The FimC-FimA (FimCA) complex was obtained by first producing FimA separately as inclusion body in the cytoplasm of *E. coli* BL21(DE3) cells as described (Puorger et al., 2011). Cells were grown in 2YT medium containing ampicillin at 100 μg/ml to OD_600_ = 1.0, gene expression was induced by adding IPTG to 1 mM and growth continued for 4 h. Cells were harvested, resuspended and lysed as described above for FimC. Inclusion bodies were prepared and solubilized essentially as described (Rudolph R, 1997). Briefly, 0.5 volumes of 60 mM EDTA-NaOH pH 7.0, 1.5 M NaCl, 6% (v/v) Triton-X-100 were added to the lysate, the suspension stirred for 30 min at 4 °C and centrifuged (30 min, 48000xg, 4 °C). Pelleted inclusion bodies were washed five times with 100 mM Tris-HCl pH 8.0, 20 mM EDTA, solubilized by suspending in 6 M guanidinium chloride (GdmCl), 50 mM Tris-HCl pH 8.0, 1 mM EDTA, 50 mM DTT (20 ml buffer per gram inclusion body), stirred for 1 h at room temperature and centrifuged (20 min, 100000xg, 20 °C). The supernatant was recovered and passed over a Superdex 200 26/60 column (GE Healthcare) equilibrated with 6 M GdmCl, 20 mM acetic acid-NaOH pH 4.0. Fractions containing FimA were pooled, diluted into 6 M GdmCl, 50 mM Tris-HCl pH 8.0 (final FimA concentration approx. 30 μM) and incubated overnight at room temperature in presence of 0.1 μM CuCl2 to allow formation of the intramolecular disulfide bond of FimA. The protein solution was first concentrated by crossflow filtration using a Vivaflow 200 system (Sartorius) and two Hydrosart cassettes (MWCO: 10 kDa) and further concentrated by ultrafiltration. For refolding and simultaneous complex formation with FimC, denatured FimA was rapidly diluted 30-fold at room temperature into 20 mM NaH_2_PO_4_-NaOH pH 7.0, 200 mM NaCl containing a ^~^2-fold molar excess of FimC and complete EDTA-free protease inhibitor cocktail (Roche). The solution was immediately desalted and the buffer exchanged by passing it over a Sephadex G25 (GE Healthcare) XK 50/20 column equilibrated with 20 mM MES-NaOH pH 5.5. For further purification, FimCA was loaded onto a Source 30S (GE Healthcare) column equilibrated with the same buffer and eluted with a linear gradient from 0-200 mM NaCl. The 1:1 complex eluted at a conductivity of approx. 7 mS/cm, appropriate fractions were pooled, concentrated by ultrafiltration and passed over a Superdex 75 26/60 column (GE Healthcare) equilibrated with 20 mM Tris-HCl (pH 8.0 at room temperature), 50 mM NaCl. Fractions containing pure FimCA complex were pooled, concentrated by ultrafiltration, aliquoted, flash-frozen in liquid nitrogen and stored at −80 °C. The identity of the proteins was confirmed by ESI-MS: Expected/measured mass for FimC was 22730.1/22729.5 Da and for FimA 15825.3/15824.5 Da.

Unfolded, disulfide-intact FimA 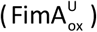 was prepared by Cu^2+^-catalyzed air oxidation of the reduced, unfolded protein as described above. The solution was then concentrated by crossflow filtration and passed over a Superdex 200 26/60 column (GE Healthcare) equilibrated in 6 M GdmCl, 20 mM Tris-HCl pH 8.0 to isolate monomeric 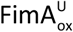. To prepare reduced, unfolded FimA, 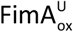 was incubated in presence of 30 mM DTT for 1 h at 37 °C. DTT was removed by desalting using a Sephadex G25 column (GE Healthcare) equilibrated in 3 M GdmCl, 20 mM Tris-HCl pH 8.0, 0.5 mM EDTA.

The FimC-FimI (FimCI) complex was prepared analogously to FimCA. *E. coli* BL21(DE3) cells carrying pfimI_cyt were used to produce FimI in form of inclusion bodies, which were isolated, solubilized and purified by size exclusion chromatography in presence of 6 M GdmCl as described above. To allow Cu^2+^-catalyzed formation of the intramolecular disulfide bond in FimI by air oxygen, the denatured protein was diluted into 6 M GdmCl, 50 mM Tris-HCl pH 8.0, 0.1 μM CuCl_2_ to a final concentration of 3 μM and incubated at room temperature overnight. After concentrating the solution by crossflow and ultrafiltration, the FimCI complex was formed by rapidly diluting denatured, disulfide-bonded FimI at 4 °C 30-fold into 20 mM acetic acid-NaOH pH 5.0 containing a ^~^2-fold molar excess of FimC and complete EDTA-free protease inhibitor cocktail (Roche). FimCI was then dialyzed against 20 mM MOPS-NaOH pH 6.7, centrifuged (10 min, 48000xg, 4 °C), loaded onto a Source 30S (GE Healthcare) column equilibrated with the same buffer and eluted with a linear gradient from 0-300 mM NaCl. Fractions containing FimCI were pooled, concentrated by ultrafiltration and passed over a Superdex 75 16/60 column (GE Healthcare) equilibrated with 20 mM Tris-HCl (pH 8.0 at room temperature), 50 mM NaCl. FimCI-containing fractions were pooled, concentrated by ultrafiltration, flash-frozen in liquid nitrogen and stored in aliquots at −80 °C. The identity of the FimCI complex was confirmed by ESI-MS: Expected/measured mass for FimC was 22730.1/22729.5 Da and for FimI 17169.3/17169.0 Da.

The FimC_His_-FimI (FimC_His_I) complex was prepared similarly. Unfolded, disulfide-intact FimI (300 μM in 6 M GdmCl, 20 mM Tris-HCl pH 8.0, 0.1 mM EDTA) was refolded by rapid, 60-fold dilution in refolding buffer (20 mM MOPS-NaOH pH 6.7) containing a small excess of FimC_His_ over FimI. The refolding mixture was dialyzed at 4 °C against 20 mM MOPS-NaOH pH 6.7 to remove residual GdmCl and applied to a Resource S column (GE Healthcare) to separate FimC_His_I from free FimC_His_. The FimC_His_I complex was eluted by applying a linear gradient from 0 - 0.3 M NaCl.

FimIt, a truncated FimI variant lacking the N-terminal donor strand, was prepared in denatured but disulfide-intact form 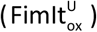 analogously as described above for FimI by using *E. coli* BL21(DE3) cells carrying pfimIt_cyt. The identity of FimIt was confirmed by ESI-MS: expected/measured mass was 15011.9/15012.5 Da.

Unfolded, disulfide-intact FimI 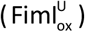 was prepared by Cu^2+^-catalyzed air oxidation of the reduced, unfolded protein as described above. The solution was then concentrated by crossflow filtration and passed over a Superdex 200 26/60 column (GE Healthcare) equilibrated in 6 M GdmCl, 20 mM Tris-HCl pH 8.0 to isolate monomeric 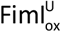. Reduced, unfolded FimI 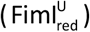 was prepared by incubating 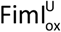 in presence of 30 mM DTT for 1 h at 37 °C. DTT was removed by desalting using a Sephadex G25 column (GE Healthcare) equilibrated in 3 M GdmCl, 20 mM Tris-HCl pH 8.0, 0.5 mM EDTA.

The FimC_His_-FimIt (FimC_His_It) complex was prepared by rapid, 60-fold dilution of 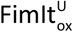 into 20 mM MOPS-NaOH pH 6.7 containing a 1.6-fold molar excess of FimC_His_. Final FimIt and FimC_His_ concentrations during refolding were 5 and 8 μM, respectively. Refolding was performed at 4 °C with stirring for 1 h. The protein solution was concentrated by crossflow filtration and dialyzed against 20 mM MOPS-NaOH pH 6.7 at 4 °C overnight. Aggregates were removed by centrifugation (20 min, 48000xg, 4° C) and the protein solution loaded onto a 6 ml Resource S column (GE Healthcare) equilibrated in 20 mM MOPS-NaOH pH 7.0. Bound FimC_His_It was eluted with a linear gradient from 0-300 mM NaCl. Fractions containing FimC_His_It were pooled and dialyzed against 10 mM Tris-HCl pH 8.0.

Oxidized DsbA (DsbA_ox_) was produced by growing *E. coli* BL21(DE3) pLysS cells transformed with plasmid pDsbA3 (Hennecke et al., 1999) at 37 °C in 2YT medium containing ampicillin at 100 μg/ml. At OD_600_ = 0.6, gene expression was induced by adding IPTG to a final concentration of 1 mM and the cells grown further for 4 h. Harvested cells were resuspended in 200 mM boric acid-NaOH pH 8.0, 160 mM NaCl, 5 mM EDTA, 1 mg/ml polymyxin B sulfate (10 ml buffer per liter of culture) and stirred for 2 h at 4 °C. After centrifugation (30 min, 48000xg, 4 °C), the supernatant was dialyzed against 10 mM MOPS-NaOH pH 7.0 overnight at 4 °C, and centrifuged again (12 min, 5000xg, 4 °C). The supernatant was loaded onto a 6 ml Resource Q column (GE Healthcare) equilibrated in 10 mM MOPS-NaOH pH 7.0. Bound DsbA was eluted with a linear 0-500 mM NaCl gradient. Fractions containing DsbA were pooled and passed over a Superdex 200 26/60 gel filtration column (GE Healthcare) equilibrated in 20 mM MOPS-NaOH, 150 mM NaCl pH 7.0. DsbA-containing fractions were pooled, dialyzed against 10 mM acetic acid-NaOH pH 4.0 overnight at 4 °C and loaded onto a 6 ml Resource S column (GE Healthcare) equilibrated in 10 mM acetic acid-NaOH pH 4.0. Bound DsbA was eluted with a linear gradient from 0-50 mM NaCl. Fractions containing DsbA were pooled, dialyzed against 20 mM MOPS-NaOH pH 7.0, flash-frozen in aliquots and stored at −20 °C.

### Preparation of ternary complexes

The ternary FimC_His_-FimI-FimAt_His_ (FimC_His_IA) and FimC_His_-FimF-FimGt (FimC_His_FG) complexes were prepared by mixing purified FimC_His_I with FimCAt_His_ or FimC_His_F with FimC_His_Gt, respectively. The ratio between the binary chaperone-subunit complexes in both cases was either 1:1, or the complex containing the truncated subunit was used at two-fold molar excess. The total protein concentration in the reactions was kept below 15 μM. After incubation for 24-36 h at 37 °C in 20 mM MOPS-NaOH pH 7.0, the ternary complexes were purified by cation exchange chromatography on a Resource S column (GE Healthcare). The FimC_His_IA complex was eluted with a linear NaCl gradient in 20 mM MOPS-NaOH pH 6.7 and eluted at ^~^200 mM NaCl. In case of FimC_His_FG, chromatography was performed in 20 mM MES-NaOH pH 5.5, and the ternary complex eluted at ^~^220 mM NaCl.

The ternary FimD_N_-FimC_His_-FimIt (FimD_N_CI) complex was prepared by mixing a 2.6-fold molar excess of purified FimD_N_ with purified FimC_His_It (final concentrations were 60 and 23 μM, respectively). After stirring the solution for 1 h at 4 °C, proteins were concentrated by ultrafiltration at 4 °C (MWCO: 10 kDa) and passed over a Superdex 75 26/60 gel filtration column (GE Healthcare) equilibrated in 20 mM sodium phosphate pH 7.4, 115 mM NaCl. Fractions containing FimD_N_CI were pooled and dialyzed against 10 mM Tris-HCl pH 8.0.

### Preparation of fluorophore-labeled FimD_N_

FimD_N_ was labeled at its N-terminus with 5/6-carboxyfluorescein succinimidyl ester as described (Nishiyama and Glockshuber, 2010). Excess label was removed and the buffer exchanged to 20 mM Tris-HCl pH 8.0 by ultrafiltration. Labeled FimD_N_ was then separated from unlabeled FimD_N_ by anion exchange chromatography and dialyzed against 20 mM Tris-HCl pH 8.0.

### Determination of protein concentrations

Protein concentrations were determined via their absorbance at 280 nm using the following molar extinction coefficients: FimDCH: 202000 M^−1^ cm^−1^; FimCF: 33015 M^−1^ cm^−1^; FimCG: 36000 M^−1^ cm^−1^; FimCA: 26680 M^−1^ cm^−1^; FimCI: 44015 M^−1^ cm^−1^; FimC_His_It: 38450 M^−1^ cm^−1^; FimC_His_Ft: 31230 M^−1^ cm^−1^; FimC_His_Gt: 35070 M^−1^ cm^−1^; FimCAt_His_: 25960 M^−1^ cm^−1^; FimC_His_FG: 44720 M^−1^ cm^−1^; FimC_His_IA: 46930 M^−1^ cm^−1^; FimD_N_CI: 44600 M^−1^ cm^−1^; FimC: 24320 M^−1^ cm^−1^; 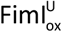: 21100 M^−1^ cm^−1^; 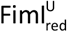: 20970 M^−1^ cm^−1^; 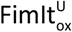: 15600 M^−1^ cm^−1^; 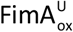: 3110 M^−1^ cm^−1^; 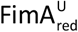: 2980 M^−1^ cm^−1^; DsbA_ox_: 23050 M^−1^ cm^−1^.

### Determination of association and dissociation rates

Rate constants for binding/dissociation of FimCI to/from FimD_N_ were determined at 23 °C in 20 mM Tris-HCl pH 8.0. Fluorescein-labeled FimDN (final concentration: 0.4 μM) was mixed with different concentrations of FimCI in a SX20 stopped-flow instrument (Applied Photophysics, UK). The reaction was monitored by recording the increase in fluorescence intensity above 515 nm (excitation at 495 nm). The fluorescence traces were globally fitted with Dynafit (Kuzmic, 1996) according to a second-order binding and first-order dissociation reaction.

The dissociation rate constant of FimC_His_ from FimC_His_I, FimC_His_IA, FimC_His_Ft or FimC_His_FG was determined as described (Vetsch et al., 2006) with minor modifications. FimC_His_I, FimC_His_IA, FimC_His_Ft or FimC_His_FG (initial concentration: 4 μM each) were incubated in presence of a 9-fold molar excess of untagged FimC in 20 mM MES-NaOH pH 6.0 at 37 °C in a shaker (300 rpm). After defined reaction times, samples were loaded at 4 °C onto a Resource S column (GE Healthcare) equilibrated in 20 mM MES-NaOH pH 6.0 (for reactions containing FimC_His_I or FimC_His_IA), 20 mM MOPS-NaOH pH 6.7 (for reactions containing FimC_His_Ft) or 20 mM MES-NaOH pH 5.5 (for reactions with FimC_His_FG), and bound proteins eluted with a linear NaCl gradient. The observed kinetics of chaperone exchange were fitted with the following single-exponential function:

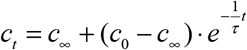

where c_∞_ is the final protein concentration at equilibrium, c_0_ the initial concentration and τ the time constant of the exchange reaction. The rate constant for FimC_His_ dissociation (k_off_) can then be calculated from the observed time constant using the known total concentrations of FimC and FimC_His_ according to:

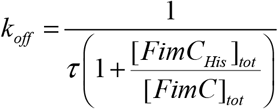

### Kinetics of pilus rod assembly

*In vitro* reconstitution of pilus assembly was performed at 23 °C (unless otherwise indicated) in 20 mM Tris-HCl pH 8.0, 50 mM NaCl, 0.05% (w/v) DDM. To convert FimD to an active FimA assembly catalyst, 0.35 μM FimDCH complex was first incubated with an 8-fold molar excess of FimCG, FimCF or both. For FimD-catalyzed assembly of FimCA both in presence and absence of FimCI, FimDCH was preincubated with both FimCG and FimCF for 30 min as this led to the highest activity of FimD with respect to FimA assembly. After preincubation, either FimCA alone, or both FimCA and FimCI were added to initiate pilus rod assembly, leading to final concentrations of 0.1 μM FimDCH, 0.8 μM FimCG, 0.8 μM FimCF, 20 μM FimCA and (if included) 0.01, 0.02, 0.04, 0.06, 0.08, 0.1, 0.14, 0.33 or 1.0 μM FimCI. Reactions were monitored by following the decrease in FimCA concentration over time. At defined assembly times, samples were loaded onto a 1 ml Resource S column (GE Healthcare) equilibrated with 20 mM MES-NaOH pH 6.0 and bound FimCA, FimCG, FimCF, FimC and FimCI eluted with a linear gradient from 0-195 mM NaCl over 29.25 ml. The absorbance at 228 nm was detected and corrected for that of a FimCA- and FimCI-free sample (in case of reactions containing both FimCA and FimCI) or for that of a FimCA-free sample (in case of reactions containing only FimCA). The areas of the FimCA and FimC peak were determined using PeakFit and the EMG function (PeakFit 4.12) and the FimCA peak area plotted against assembly time. Using Dynafit (Kuzmic, 1996), all FimD-catalyzed kinetics were fitted numerically according the mechanism depicted in Figure 2D, with the rate constants for uncatalyzed FimA assembly (k_(A-A)uncat_), binding of FimCA and FimCI to FimD_N_ (k_on_) and dissociation of FimCA and FimCI from FimD_N_ (k_off_) fixed to k_(A-A)uncat_ = 0.73 M^−1^ s^−1^, k_on_ (FimCA) = 1.1·10^7^ M^−1^ s^−1^, k_on_ (FimCI) = 1.45 · 10^8^ M^−1^ s^−1^, k_off_ (FimCA) = 267 s^−1^ and k_off_ (FimCI) = 94 s^−1^, respectively. Each kinetic trace was first fitted individually with the initial FimCA concentration fixed to 20 μM, the initial concentration of activated FimDCH molecules fixed to 0.1 μM, and the FimCA response value and DSE rate constants k_(A-F)_, k_(A-A)_ and k_(I-A)_ as fitting parameters. For normalization and conversion of FimCA peak areas to FimCA concentrations, experimentally determined FimCA peak areas of each kinetic trace were then divided by the corresponding fitted response value. The normalized data were then fitted globally, now with the FimCA response value fixed to 1, the initial FimCA concentration fixed to 20 μM and the concentration of activated FimDCH molecules and k_(A-F)_, k_(A-A)_ and k_(I-A)_ as fitting parameters.

Uncatalyzed FimCA assembly was measured similarly by i) omitting FimDCH from the reaction mix and using 1 μM FimCI, the highest FimCI concentration used in the catalyzed assembly reactions or ii) incubating 20 μM FimCA alone. Both sets of raw data were first individually fitted according to an irreversible, second-order dimerization reaction, normalized using the fit value for the peak area at t = 0 and the known initial FimCA concentration, and then fitted globally with shared k_(A-A)uncat_. Samples for negative-stain electron microscopy were removed after 65 min of assembly, flash-frozen in liquid nitrogen and stored at −20 °C for later analysis.

### Electron microscopy and pilus length measurements

Samples for negative-stain electron microscopy were prepared essentially as described (Hahn et al., 2002). *E. coli* strains W3110, W3110Δ*fimA* and W3110Δ*fimA* [pCG1-AC] were grown statically at 37 °C in 2YT medium (supplemented with ampicillin at 100 μg/ml in case of W3110Δ*fimA* [pCG1-AC]). *In vivo*, type 1 pili are assembled within minutes during the exponential growth phase of the culture but assembly slows considerably when the culture approaches the stationary phase (Dodd and Eisenstein, 1984). Therefore, in case of W3110Δ*fimA* [pCG1-AC], anhydrotetracycline (aTc, Sigma) was added to the growth medium just prior to inoculation to final concentrations of 0, 1, 2, 5, 7.5, 10, 12.5, 15 and 17.5 ng/ml to ensure that FimA and FimC expressed from pCG1-AC would be available for assembly, in particular during the exponential phase of the bacterial growth curve. After 10 h of growth, 2 ml of cell culture were centrifuged (10 min, 2500xg, 4 °C), the pelleted cells resuspended in 100 μl NaH_2_PO_4_-NaOH pH 7.5 and a 3 μl drop adsorbed for 20 s to a carbon-coated copper grid (300 mesh, Quantifoil) that had been glow-discharged for 30 s at a current of 25 mA. Excess liquid was removed by blotting the grid with filter paper, followed by negatively staining the sample for 30 s with a 20 μl drop of 1% (w/v) phosphotungstic acid-NaOH pH 7.6. After removing excess stain with filter paper, the grid was air-dried and images recorded using a Morgagni 268 microscope (FEI) operated at an acceleration voltage of 100 kV and equipped with a 1376×1032 pixel CCD camera. Cell-bound pili of W3110 and W3110Δ*fimA* [pCG1-AC] cells were released from the outer membrane by incubating the remainder of the cell suspension in a water bath at 90 °C for 10 min. After centrifugation (10 min, 2500xg, room temperature), the supernatant was used for grid preparation as above. For samples produced by *in vitro* pilus assembly reactions, electron microscopy grids were prepared as above but using 2% (w/v) phosphotungstic acid-NaOH pH 7.2 for staining. To investigate a larger area of the grid and thus avoid bias towards shorter pili, an array of 5×5 overlapping micrographs was acquired using the Multiple Image Acquisition function of the Morgagni user interface. These 25 micrographs were then stitched using the Grid/Collection stitching plugin of Fiji (Preibisch et al., 2009; Schindelin et al., 2012). In case of W3110Δ*fimA* [pCG1-AC] grown in presence of 17.5 ng/ml aTc, the investigated grid area was further increased by manually superimposing four overlapping 5×5 arrays with CorelDRAW (Corel Corporation). The lengths of 200 pili per sample were measured using TrakEM2 (Cardona et al., 2012). Data were binned, analyzed and plotted using OriginPro 9.1 (OriginLab).

### Kinetics of DsbA-catalyzed oxidation and FimC-catalyzed folding of FimA and FimI

Rate constants for DsbA-catalyzed oxidation of reduced, unfolded FimI and FimA 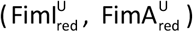 were determined at 25 °C and pH 8.0 by 10:1 mixing of oxidized DsbA (5.45 μM in 20 mM Tris-HCl pH 8.0) with 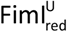 (8.25 μM in 3 M GdmCl, 20 mM Tris-HCl pH 8.0) or 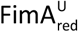 (8.25 μM in 3 M GdmCl, 20 mM Tris-HCl pH 8.0) using a SX18.MV stopped-flow mixing instrument (Applied Photophysics, UK). Initial concentrations were 5 μM oxidized DsbA and 0.75 μM 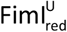 or 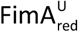. Reactions were monitored by recording the change in intrinsic fluorescence intensity above 320 nm (excitation at 280 nm). For oxidation of 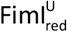, fluorescence traces were fitted according to a pseudo-first order reaction (6.7-fold excess of oxidized DsbA over 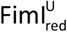) or a second-order reaction with identical initial concentrations (0.75 μM of both oxidized DsbA and 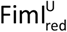), and then normalized. Normalized data were globally fitted according to an irreversible, second-order reaction and sharing the rate constant for oxidation and the initial 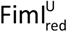 concentration among the two datasets.

Rate constants for FimC-catalyzed folding of oxidized, unfolded FimI and FimA 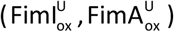 were determined by 10:1 mixing of FimC (5.45 μM in 20 mM Tris-HCl pH 8.0) with 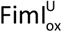 or 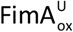 (8.25 μM in 3 M GdmCl, 20 mM Tris-HCl pH 8.0) using stopped-flow fluorescence spectroscopy as above and fitting the recorded traces according to a pseudo-first order reaction.

### Real-time PCR

*E. coli* strains W3110, W3110Δ*fimA* and W3110Δ*fimI* were grown overnight in LB medium at 37 °C with shaking. RNA was extracted using the RNeasy Mini Kit (Qiagen). Residual DNA was removed by on-column DNase digestion. Total yield of purified RNA was determined by absorbance spectroscopy and extracted RNA diluted to a final concentration of 1 μg/μl. cDNA synthesis was carried out using 1 μg of RNA and the High Capacity cDNA Reverse Transcription Kit (Applied Biosystems) according to manufacturer’s instructions. Real-time PCR was performed using TaqMan gene expression master mix and TaqMan gene expression assays (Applied Biosystems) on an ABI 7900 instrument with the following cycling parameters: 1) 50 °C for 2 min; 2) 95 °C for 10 min; 3) 95 °C for 15 sec and 60 °C for 1 min, repeated 40 times. Primers used were: for *fimI*: fimI_FW377 (5′-ATGAAGGAAACCTCGTACCG-3′), fimI_RV451 (5′-CGATATTTGGCGATGAAATG-3′) and fimI_probe406 (5′-CCTCCAGCAAACTGGAAACGGC-3′); for *fimA*: fimA_FW134 (5′-CAGTTGATGCAGGCTCTGTT-3′), fimA_RV251 (5′-AGATGCAACATTGGTATCGC-3′) and fimA_probe 2 (5′-CCTTCCTGTGCCAGCGATGC-3′); for *fimC*: fimC_FW342 (5′-CCGGGAAAGTTTATTCTGGA-3′), fimC_RV450 (5′-CTAATTTAGCCGGGCGATAG-3′) and fimC_probe (5′-CAGCTCGCAATTATCAGCCGCA-3′); for GAPDH: GAPDH_FW (5′-AGCTGCAACTTACGAGCAGA-3′), GAPDH_RV (5′-CTTTAGCATCGAACACGGAA-3′) and GAPDH_probe (5′-TTCGGTGTAGCCCAGAACGCC-3′). All probe primers were labelled with FAM and TAMRA at their 5′- and 3′-end, respectively. PCRs were set up in triplicate and repeated three times. Data were analyzed with the SDS 2.3 software. Raw C_T_ values were transformed to relative transcript levels by the 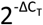 method using GAPDH as internal reference gene (Schmittgen and Livak, 2008).

### Yeast agglutination assay

Yeast agglutination assays were performed as described (Vetsch M, 2005) with minor modifications. *E. coli* strains W3110, W3110Δ*fimA*, W3110Δ*fimI* and W3110Δ*fimI* [pKTCTET-IC] were grown at 37 °C in LB medium (supplemented with ampicillin at 100 μg/ml in case of W3110Δ*fimI* [pKTCTET-IC]) under static conditions or with shaking at 210 rpm in a Multitron shaker (Infors HT). After 14.5 h of growth, cells were harvested by centrifugation (10 min 4000xg, 4 °C), resuspended in 20 mM NaH_2_PO_4_-NaOH pH 7.0, 150 mM NaCl and diluted to OD_600_ = 1.0 with the same buffer. In 24-well Linbro plates, 250 μl of the bacterial suspension were then mixed with 50 μl of a 10% (w/v) suspension of dry baker’s yeast prepared in 20 mM NaH_2_PO_4_-NaOH pH 7.0, 150 mM NaCl and imaged after 5 min of incubation. To test for mannose specificity, bacteria were shortly preincubated with 1.2 M D-mannose or D-glucose prior to mixing with the yeast cells.

### Monte Carlo simulations

Monte Carlo simulations of pilus rod assembly reactions in absence and presence of FimCI were performed using MATLAB and its accessory application SimBiology (MATLAB R2017a). To be able to assess the number of FimA monomers incorporated by a given FimD molecule at the end of the simulation, the mechanism depicted in Figure 2D was modified such that a separate pool for each polymeric and intermediate species was defined (see Figure S7 for a schematic of the first three steps of the polymerization reaction). The maximum possible number of FimA monomers per pilus rod was set to 2200 to ensure inclusion of potentially very long pili. Simulations were run with initial numbers of 2000 activated FimDCH, 487805 FimCA and, for reactions involving FimI, 244, 488, 976, 1463, 1951, 2439, 3415 or 8049 FimCI molecules. Using a concentration of 0.082 μM for activated FimDCH (the value obtained from the global fit of the experimental kinetic data) this corresponded to 20 μM FimCA, 0.01, 0.02, 0.04, 0.06, 0.08, 0.1, 0.14 or 0.33 μM FimCI, and a reaction volume of 4.05 · 10^−14^ l. The rate constants used were k_on_ = 1.1 · 10^7^ M^−1^ s^−1^ and k_off_ = 267 s^−1^ for binding/dissociation of FimCA to/from activated FimDCH; k_on_ = 1.45 · 10^8^ M^−1^ s^−1^ and k_off_ = 94 s^−1^ for binding/dissociation of FimCI to/from activated FimDCH; k_(A-F)_ = 5.5 · 10^-3^ s^−1^ for incorporation of the first FimA monomer when FimF is at the growing end of the pilus; k_(A-A)_ = 1.4 s^−1^ for incorporation of FimA monomers when FimA is at the growing end of the pilus; k_(I-A)_ = 0.14 s^−1^ for incorporation of FimI; and k_(A-A)uncat_ = 0.73 M^−1^ s^−1^ for uncatalyzed polymerization of FimCA. Second-order rate constants were converted to their stochastic values by dividing by N_A_·V (for binding of FimCA or FimCI to activated FimDCH) or by multiplying by 2/(N_A_·V) (for the uncatalyzed polymerization of FimCA), where N_A_ is Avogadro’s constant and V the reaction volume (Wilkinson, 2012). All rate constants were assumed to be independent of polymer length. The simulations were run for 65 minutes using the stochastic simulation algorithm (Gillespie, 1977) as solver and LogDecimation set to 10000. Pilus lengths were calculated by multiplying the number of FimA monomers assembled by each FimD molecule by 0.8 nm, the axial rise of the type 1 pilus rod (Hospenthal et al., 2017). The complete script used for the simulations is provided at the end of the supplemental information. Pili shorter than 15 nm were removed from the data sets and thus excluded from the comparison with the experimental length distributions as it was not possible to distinguish such short pili (if present) from background in the EM micrographs.

### Crystallization, data collection and structure determination

Crystals of the FimC_His_-FimIt complex were obtained at 4 °C via sitting drop vapor diffusion by mixing 1.5 μl of protein solution (22 mg/ml in 10 mM Tris-HCl pH 8.0) with 1 μl of precipitant solution containing 4.5 M sodium formate, 0.1 M sodium cacodylate at pH 6.5. First crystals appeared after three days. Crystals were cryo-protected by adding 8 μl of 20% ethylene glycol in mother liquor solution directly to the drop and then flash-cooled in liquid nitrogen.

Crystals of the FimD_N_-FimC_His_-FimIt complex were obtained at 4 °C via sitting drop vapor diffusion by mixing 1 μl of protein solution (19.5 mg/ml in 10 mM Tris-HCl pH 8.0) with 1 μl of precipitant solution containing 0.1 M Hepes pH 8.4, 15 % PEG 4000. Crystals were cryo-protected in mother liquor containing 30% (v/v) ethylene glycol and flash-cooled in liquid nitrogen.

Diffraction data for FimC_His_-FimIt and FimD_N_-FimC_His_-FimIt were collected at 100 K using a wavelength of 1 Å at beamline X06DA (Paul Scherrer Institute, Villigen, Switzerland) on a Pilatus 2MF pixel detector. For FimC_His_-FimIt, data were processed to a final resolution of 1.75 Å with XDS (Kabsch, 2010) in space group P 31 2 1 with one complex per asymmetric unit. For FimD_N_-FimC_His_-FimIt, data were processed with XDS to a final resolution of 1.70 Å in space group P 21 21 21. The structures were solved by molecular replacement using PHASER (McCoy et al., 2007). For FimC_His_-FimIt, the structure of the binary FimC-FimAt complex (pdb code 4DWH) (Crespo et al., 2012) was used as a search model. For FimD_N_-FimC_His_-FimIt, the structure of the FimC_His_-FimIt complex was used as a search model. After molecular replacement, initial refinement revealed the presence of FimD_N_, which was then copied from PDB 1ze3. Refinements were performed with PHENIX (Liebschner et al., 2019), and COOT (Emsley et al., 2010) was used for manual rebuilding of the model. Due to missing electron density, the following residues were omitted from the final models: for FimC_His_-FimIt: FimI residues 123-127; for FimD_N_-FimC_His_-FimIt: FimD_N_ residues 11-14, 124-125 and FimI residues 119-121.

Crystals of the FimC_His_-FimI-FimAt_His_ complex were obtained at 293 K via the vapor diffusion sitting drop method by mixing 2 μl of protein solution (10.2 mg/ml, containing less than 20 mM NaCl) with 2 μl of precipitant solution containing 0.8 M sodium formate, 100 mM Tris/acetic acid pH 8.5 and 16% PEG 4000 (w/v). Crystals grew to full size within 8 days. Diffraction data to 2.8 Å resolution were collected at 100 K on a Mar225 detector at beamline X06SA of the Swiss Light Source (PSI Villigen, Switzerland). For cryoprotection, a solution containing 16% PEG4000, 0.8 M sodium formate, 100 mM Tris-acetic acid pH 8.5, and 22% (v/v) ethylene glycol was used. Crystals were bathed in the cryo solution for 30-60 seconds before direct flash cooling in the cryostream. Data were processed with XDS (Kabsch, 2010) in space group P6(1). The structure was solved by molecular replacement with PHASER (McCoy et al., 2007), using the structure of the FimC-FimAt complex (PDB file 3SQB) (Crespo et al., 2012) as search model for FimC_His_-FimI, and FimAt from the same complex as search model for the FimAt subunit in FimC_His_-FimI-FimAt_His_. Based on Matthews volume considerations the asymmetric unit was expected to contain at least two copies of the ternary complex. The PHASER runs could locate a complete copy of FimC_His_-FimI-FimAt_His_ and an additional FimC_His_-FimI complex. The lacking FimAt molecule in the second copy of the ternary complex was placed manually by superimposing the first FimC_His_-FimI-FimAt_His_ complex onto the second FimC_His_-FimI. The second FimAt molecule was found to fit into the electron density calculated from the molecules already located by PHASER. No further complexes could be located and we concluded that there are two FimC_His_-FimI-FimAt_His_ complexes in the asymmetric unit, corresponding to a Matthew coefficient of 4.64 A^3^/Da (solvent content of 73.5%). This quite high solvent content is readily explained by the presence of large channels in the crystal packing. The structure was refined by successive rounds of manual model building with COOT (Emsley et al., 2010) and refinement with PHENIX (Liebschner et al., 2019) using TLS groups defined by PHENIX and scale factors for the X-ray/stereochemistry weight (wxc_scale) and X-ray/ADP weight (wxu_scale) fixed to 0.25 and 1.66, respectively. Due to missing electron density, the following residues were omitted from the final model: residues 1-10 and 124-126 (chain B, FimI), residues 14-24, 57-69, 121-129 and 159 (chain C, FimAt), residues 1-5 and 123-127 (chain E, FimI) and residues 14-17 (chain F, FimAt). Atomic coordinates were deposited in the Protein Data Bank (PDB) with accession codes 6SWH (FimC_His_IA), 7B0W (FimC_His_It) and 7B0X (FimD_N_CI).

Validation with MolProbity (Chen et al., 2010) showed that all final models possessed good stereochemical quality. Data collection and refinement statistics are shown in Table S2. Structural superpositions of proteins and Cα RMSD values were calculated using SUPERPOSE and secondary-structure matching (Krissinel and Henrick, 2004). The interface area between FimI and FimAt in the FimC_His_IA complex was calculated using PISA (Krissinel and Henrick, 2007), all structural representations were created with PyMOL (Schrodinger, 2015). The structure-based sequence alignment of all type 1 pilus subunits was created with Expresso (Armougom et al., 2006), using both SAP and TMalign for structural alignment, and rendered using ESPript 3.0 (Robert and Gouet, 2014). Secondary structure of FimI was assigned using DSSP (Kabsch and Sander, 1983; Touw et al., 2015).

## SUPPLEMENTAL INFORMATION

**Figure S1.**
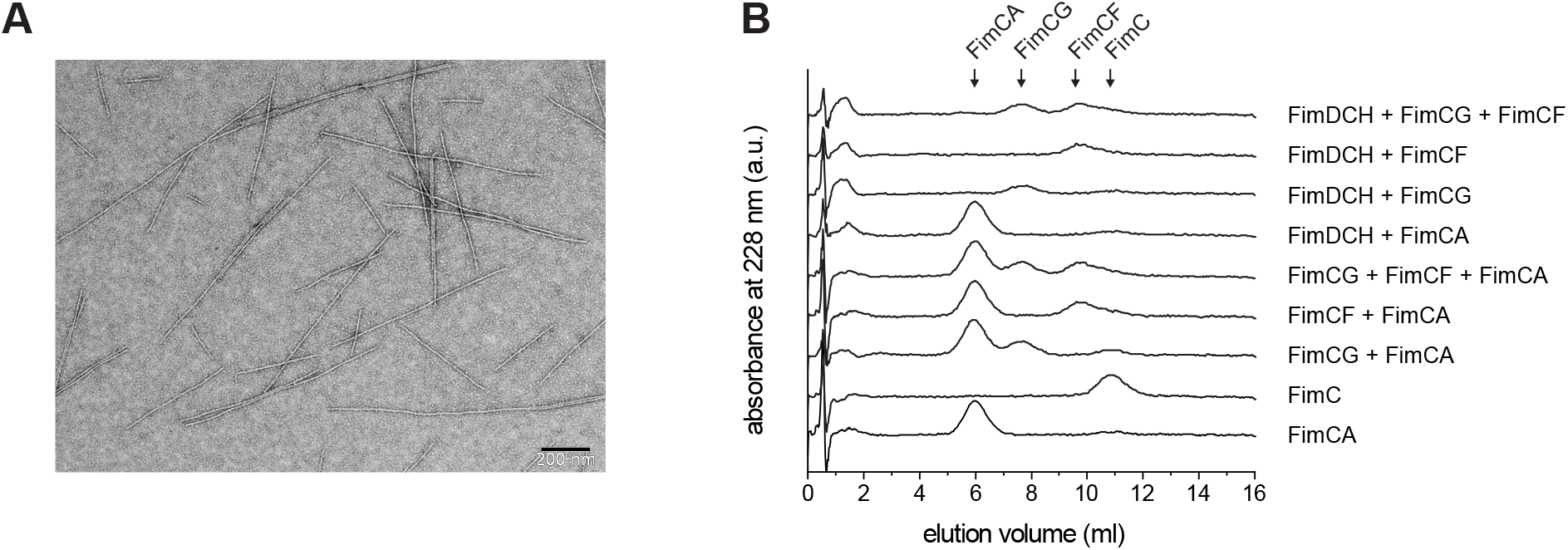
Control experiments regarding the reconstitution of type 1 pilus assembly, Related to Figure 1. (A) Negative-stain electron micrograph of pilus rods assembled *in vitro* by FimD. FimDCH (0.2 μM) was incubated with a 20-fold molar excess of FimCF for 5 min at 37 °C. FimCA was added to a final concentration of 10 μM resulting in a 100-fold excess over FimD, and the reaction incubated further until completion (scale bar = 200 nm). (B) Cation exchange chromatography profiles of control reactions at 37 °C proved that changes in FimCA and FimC peak intensity reflected FimD-catalyzed pilus rod assembly only and were not caused by other reactions, for instance between FimCA and FimCG/FimCF. Reactions including FimDCH and FimCG/FimCF were first preincubated for 5 min before addition of buffer and additional incubation for 10 min. All other reactions were analyzed after 10 min of incubation. Concentrations were 0.25 μM FimDCH, 2 μM FimCG and/or FimCF, 5 μM FimCA and 5 μM FimC.

**Figure S2.**
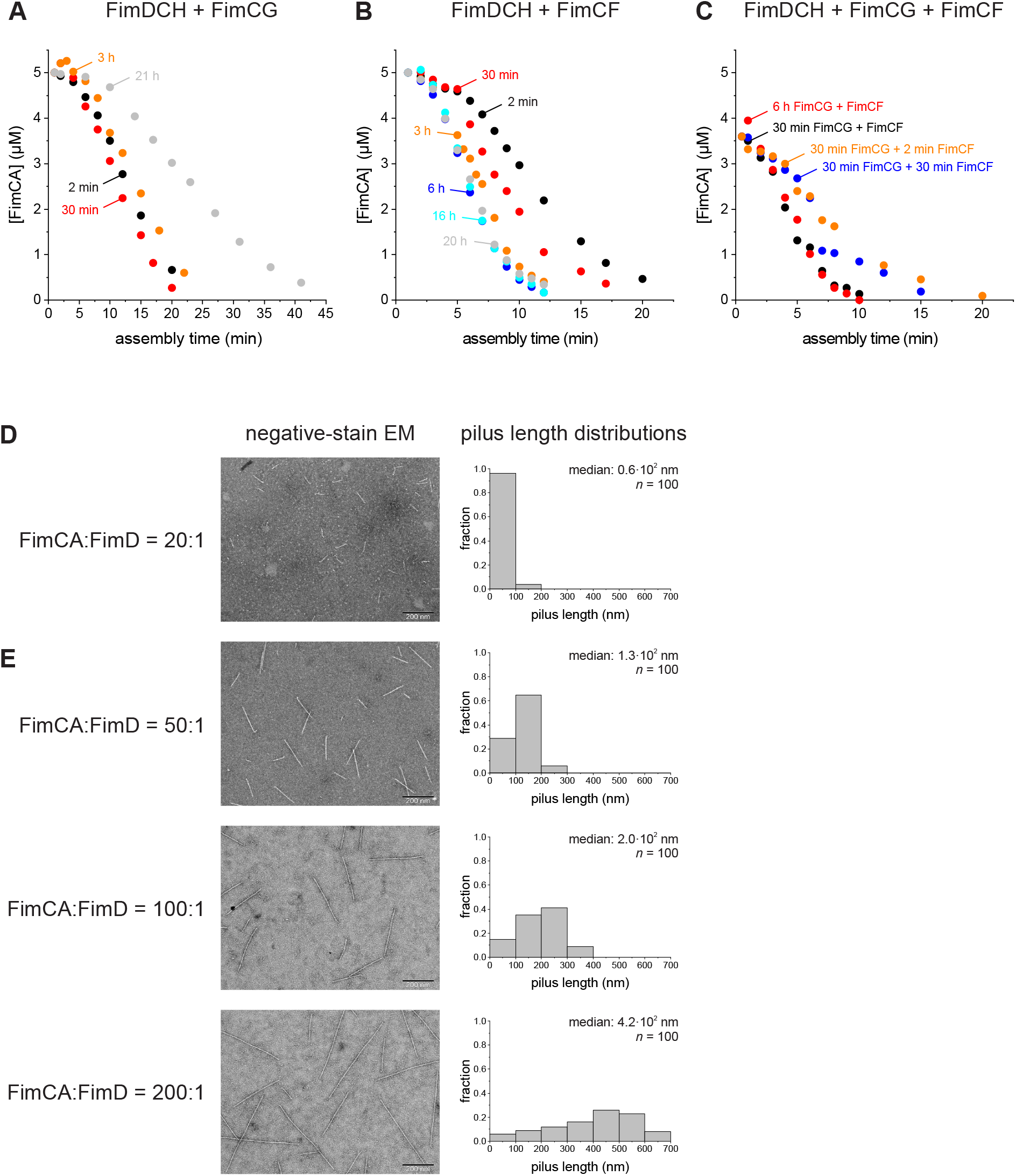
Optimization of reaction conditions for type 1 pilus rod assembly *in vitro* at pH 8.0 and 23 °C, Related to Figure 1 and Figure 2. (A) Kinetics of FimD-catalyzed FimA assembly measured after activation of 0.35 μM FimDCH by 2.8 μM FimCG for the indicated periods of time. Initial concentrations of FimDCH, FimCG and FimCA in the assembly reaction were 0.25, 2 and 5 μM, respectively. (B) Kinetics of FimD-catalyzed FimA assembly measured after activation of 0.35 μM FimDCH by 2.8 μM FimCF for the indicated periods of time. Initial concentrations of FimDCH, FimCF and FimCA in the assembly reaction were 0.25, 2 and 5 μM, respectively. (C) Kinetics of FimD-catalyzed FimA assembly measured after activation of FimDCH by sequential addition of FimCG and FimCF (shown in orange and blue) or by simultaneous co-incubation of all three complexes (shown in black and red) for the indicated periods of time. For sequential activation, FimCG and FimCF were used at 2.8 μM each, corresponding to an 8-fold excess of FimCG over FimDCH in the first, and a 14-fold excess of FimCF over FimDCH in the second step of the reaction. Initial concentrations of FimDCH, FimCG, FimCF and FimCA in the pilus rod assembly reaction were 0.18, 1.4, 2.5 and 3.6 μM, respectively. For simultaneous FimDCH activation, FimDCH, FimCG and FimCF were used at 0.35 and 2.8 μM, respectively. Initial concentrations of FimDCH, FimCG, FimCF and FimCA in the pilus rod assembly reaction were 0.18, 1.4, 1.4 and 3.6 μM, respectively. Kinetics in (A), (B) and (C) were normalized using the known initial FimCA concentration and the peak area of the first data point. (D and E) Negative-stain electron micrographs and length distributions of pilus rods assembled using a 20-, 50-, 100- or 200-fold excess of FimCA over FimD. FimDCH (0.35 μM) was activated by preincubation with an 8-fold excess of both FimCG and FimCF for 30 min. FimCA was added to 3.6, 9, 18 or 36 μM and samples analyzed after 20 min of assembly (scale bar = 200 nm). For pilus length distributions, the lengths of 100 pili were determined.

**Figure S3.**
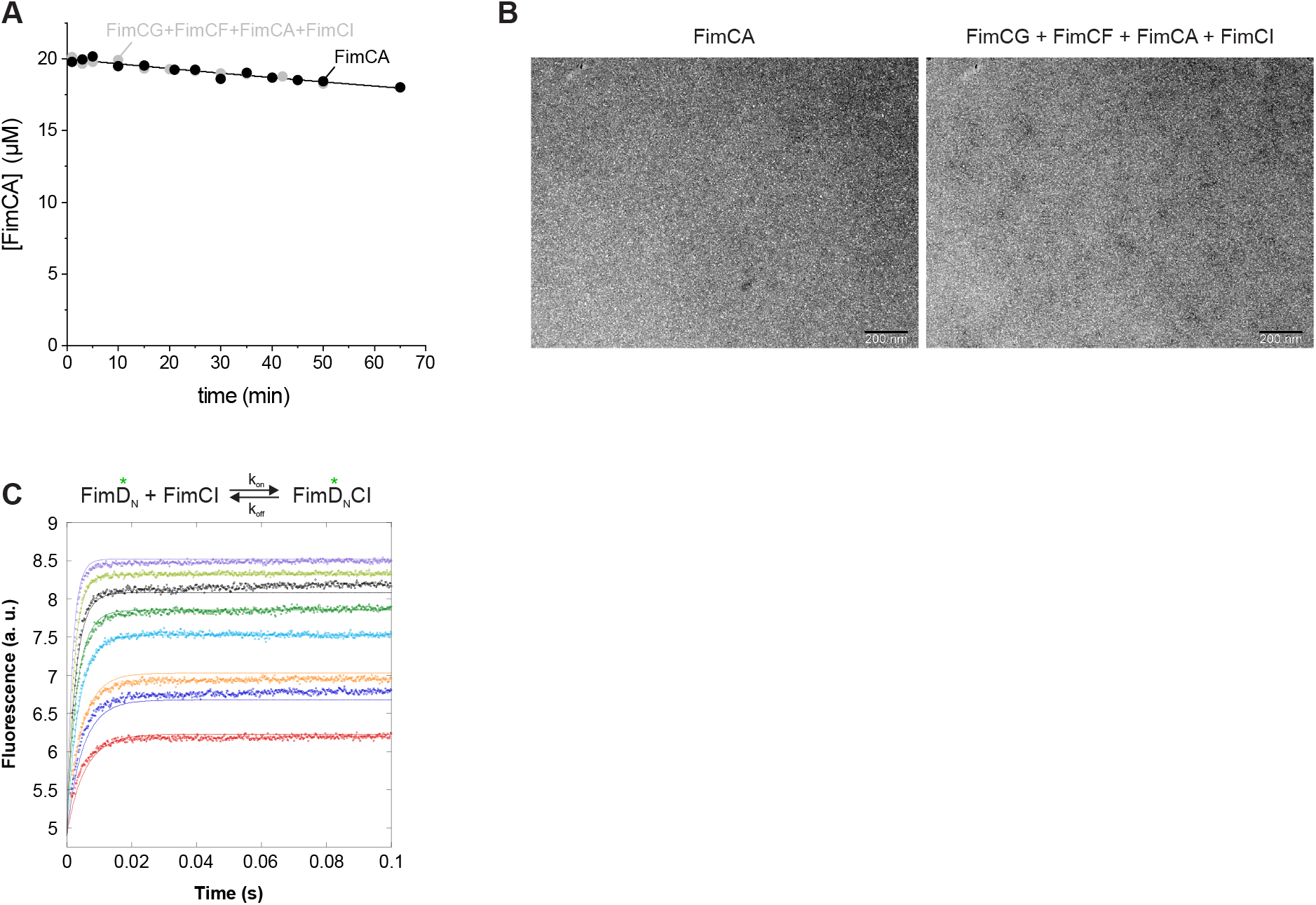
Kinetics of uncatalyzed FimA assembly and binding of FimCI to FimD_N_, Related to Figure 2. (A) Kinetics of uncatalyzed FimA assembly. FimCA (20 μM) was either incubated alone or in presence of 0.8 μM FimCG, 0.8 μM FimCF and 1 μM FimCI for 65 min at 23 °C. Both data sets were globally fitted according to an irreversible second-order dimerization reaction (solid line) yielding k_(A-A)uncat_ = 0.73 ± 0.02 M^−1^ s^−1^. (B) Negative-stain electron micrographs of samples from the uncatalyzed FimA assembly reactions shown in (A), taken after 65 min of assembly (scale bar = 200 nm). (C) Binding kinetics for fluorescently labeled FimDN and FimCI measured by stopped-flow fluorescence spectroscopy at pH 8.0 and 23 °C (green asterisks indicate the fluorophore). Initial concentrations were 0.4 μM FimD_N_ and 0.4, 0.6, 0.8, 1.2, 1.6, 2.0, 2.6 and 3.4 μM FimCI. The solid lines represent a global fit of the data according to a second-order binding and first-order dissociation reaction yielding k_on_ = 1.45 · 10^8^ M^−1^ s^−1^ and k_off_ = 94 s^−1^.

**Figure S4.**
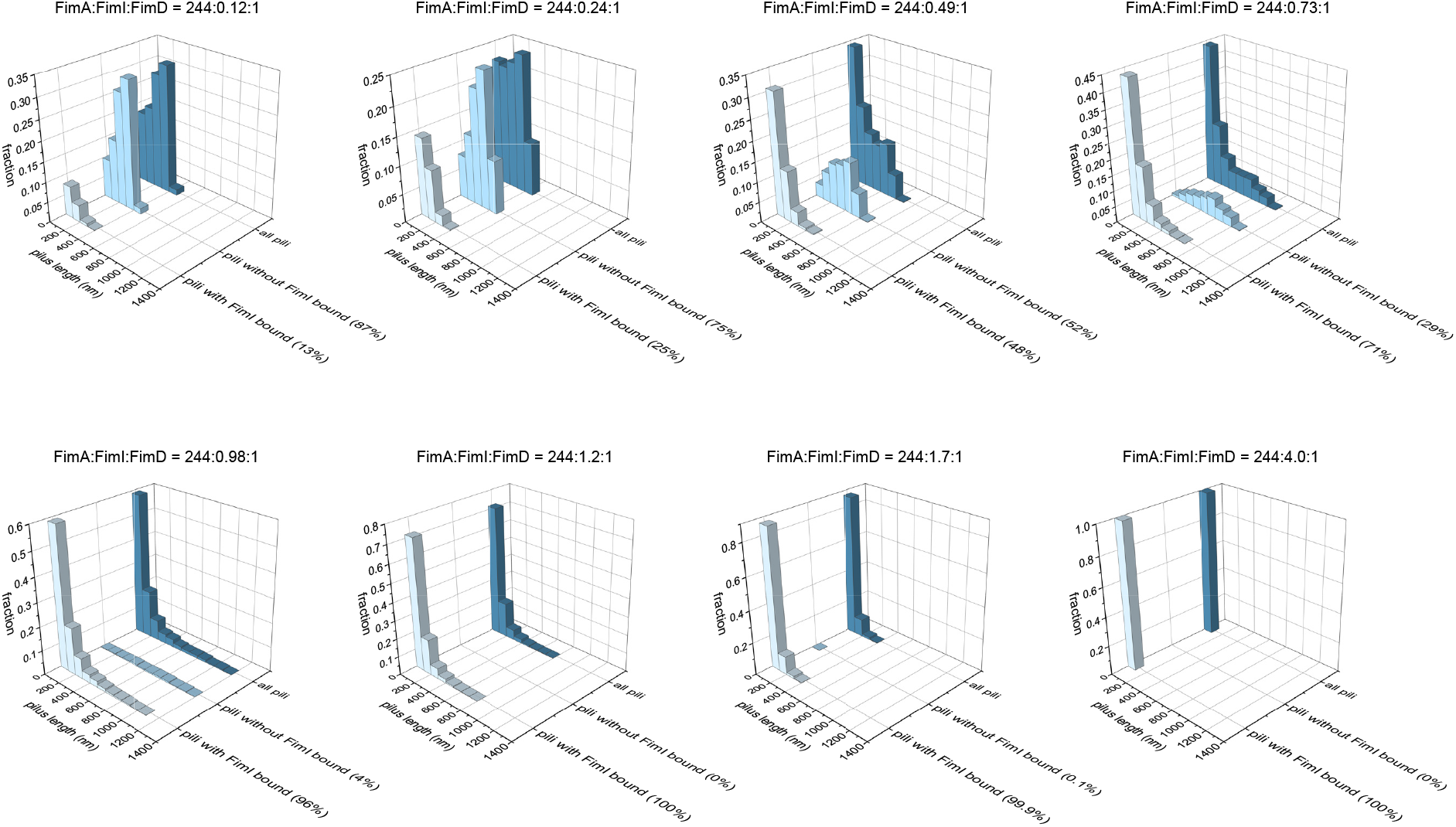
FimI-terminated pili are predominantly short, Related to Figure 3. Pilus length distributions obtained after simulating type 1 pilus rod assembly in the presence of different FimCI concentrations. Each panel shows the distribution comprising all pili (in blue) together with the two underlying distributions of FimI-terminated pilus rods (in gray) and non-terminated rods (in light-blue). For each simulation, the fractions of FimI-terminated and non-terminated pilus rods are given in parentheses. Ratios between FimCA, FimCI and FimD are indicated above each panel.

**Figure S5.**
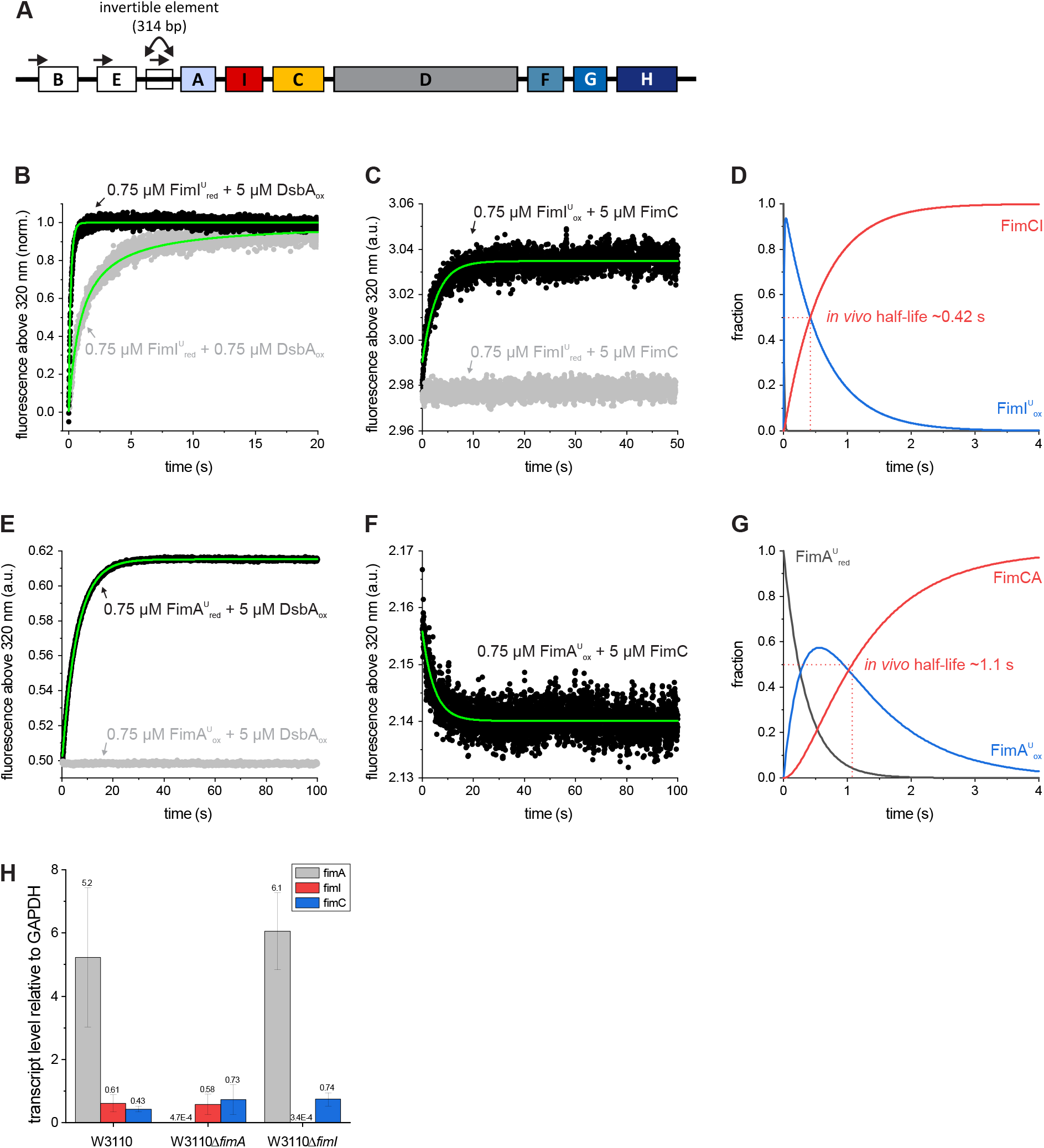
Type 1 pilus gene cluster, kinetics of oxidation and folding of FimI and FimA measured by stopped-flow fluorescence spectroscopy at pH 8.0 and 25 °C and transcript levels of *fimA*, *fimI* and *fimC*, Related to Figure 3. (A) Schematic representation of the type 1 pilus gene cluster (Schwan, 2011). Promoter regions are indicated by horizontal arrows. Expression of the *fimA*, *fimI*, *fimC*, *fimD*, *fimF*, *fimG* and *fimH* genes is controlled by a promoter located upstream of *fimA* within a 314 bp-long invertible DNA element (Abraham et al., 1985; Olsen and Klemm, 1994). The orientation of this element can be switched by two recombinases, FimB and FimE, encoded upstream of the element(Klemm, 1986). (B) Kinetics of DsbA-catalyzed disulfide bond formation in reduced, unfolded FimI 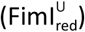 recorded using identical initial concentrations of oxidized DsbA (DsbA_ox_) and 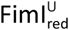 (0.75 μM each) or under pseudo-first order conditions with a 6.7-fold excess of DsbA_ox_ over 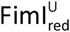. After normalization, both datasets were globally fitted according to an irreversible, second-order reaction (green line). The rate constant for oxidation and the initial 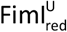 concentration were shared among the datasets. (C) The kinetic of FimC-catalyzed folding of oxidized, unfolded FimI (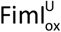, 0.75 μM) was recorded under pseudo-first order conditions (shown in black). The solid line corresponds to a fit according to a single-exponential function. FimC specifically recognized disulfide-intact, unfolded FimI as no fluorescence change was detected when reduced, unfolded FimI was mixed with FimC (shown in gray). (D) Simulated kinetics for oxidative folding of FimI *in vivo*. The fractions of 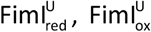 and native FimCI complex were calculated using the rate constants for DsbA-catalyzed oxidation and FimC-catalyzed folding determined in (B) and (C), the experimentally determined periplasmic concentrations of DsbA (86 μM) and FimC (23 μM) and assuming pseudo-first order conditions for FimI, i.e. that DsbA and FimC are in excess over 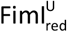 and 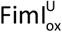 in the periplasm, respectively. The resulting, theoretical *in vivo* half-life for formation of the native FimCI complex is indicated. (E) Kinetic of DsbA-catalyzed disulfide bond formation in reduced, unfolded FimA (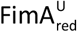, 0.75 μM) recorded under pseudo-first order conditions (shown in black). The solid line corresponds to a fit according to a single-exponential function. No reaction occurred when oxidized, unfolded FimA (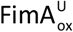) was mixed with oxidized DsbA (shown in gray). (F) Kinetic of FimC-catalyzed folding of 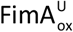 (0.75 μM) recorded under pseudo-first order conditions and fitted according to a single-exponential function (solid line). (G) Simulated kinetics for oxidative folding of FimA *in vivo*. The fractions of 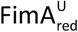, 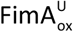 and native FimCA complex were calculated using the rate constants for DsbA-catalyzed oxidation and FimC-catalyzed folding determined in (E) and (F), the experimentally determined periplasmic concentrations of DsbA (86 μM) and FimC (23 μM) and assuming pseudo-first order conditions for FimA. The resulting, theoretical *in vivo* half-life for formation of the native FimCA complex is indicated. (H) Transcript levels of *fimA*, *fimI* and *fimC* in *E. coli* W3110, W3110Δ*fimA* and W3110Δ*fimI* cells determined by real-time PCR relative to the transcript level of GAPDH. Data are means ± s.e. (*n* = 3).

**Figure S6.**
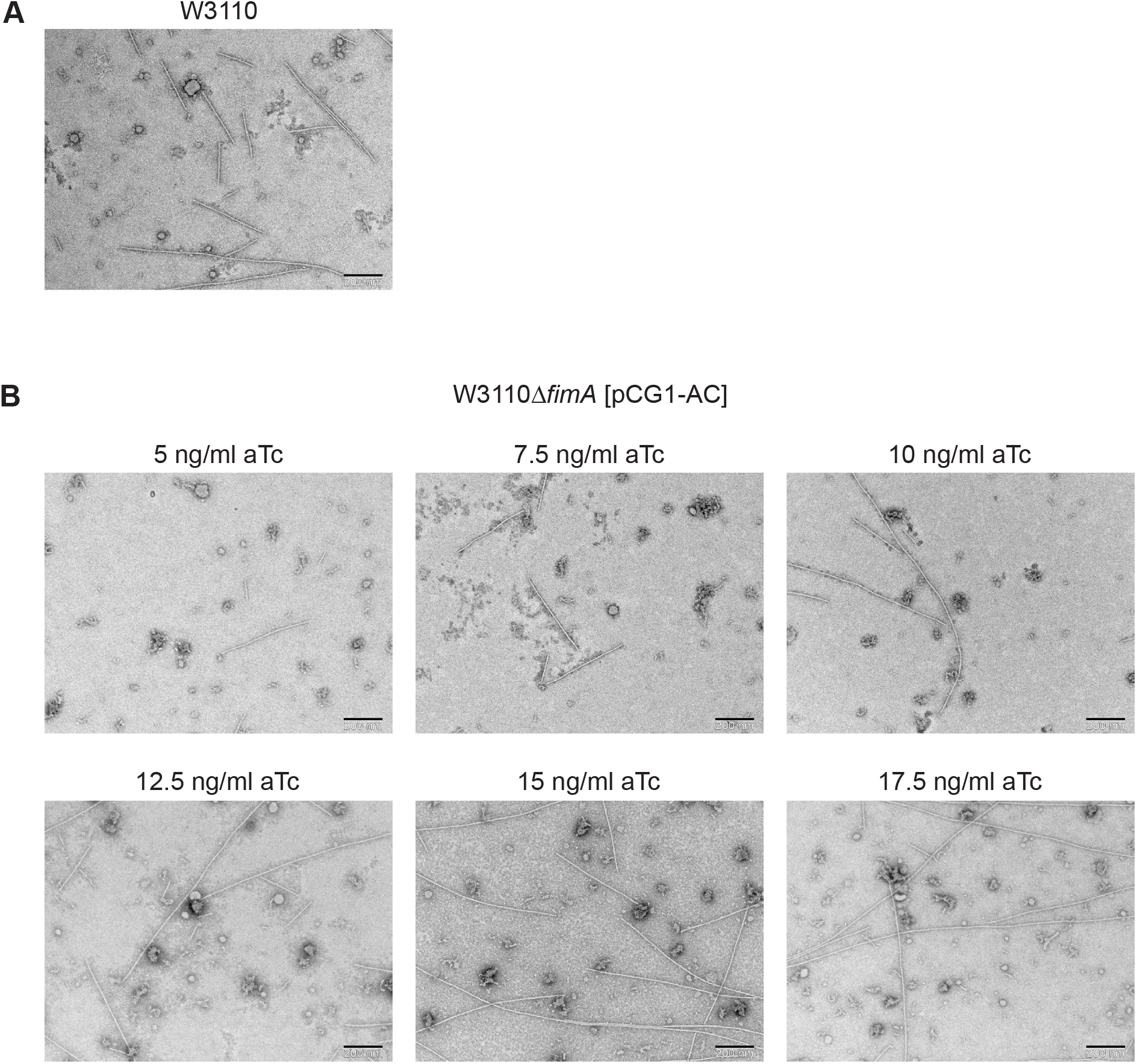
Negative-stain electron micrographs of pili released from cells, Related to Figure 4. Pili released from (A) *E. coli* W3110 cells or (B) W3110Δ*fimA* cells harboring pCG1-AC and grown in presence of the indicated anhydrotetracycline (aTc) concentrations. Scale bar = 200 nm.

**Figure S7.**
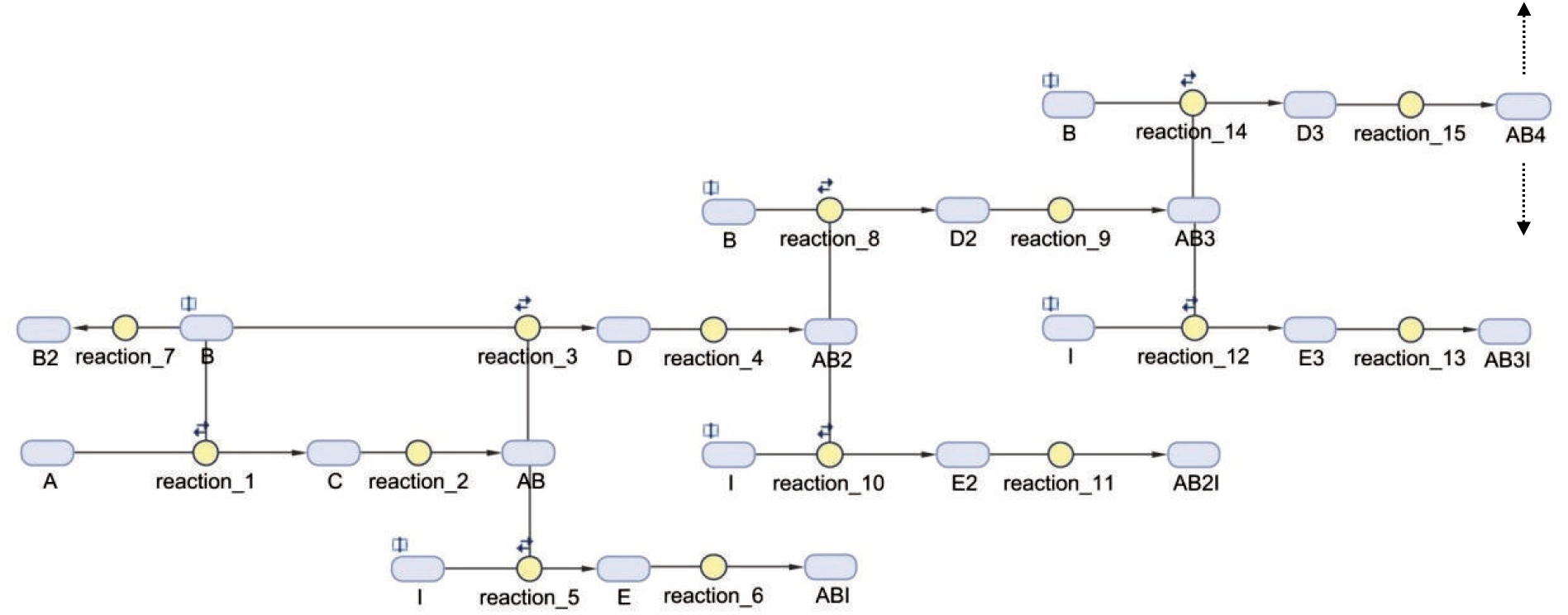
Stochastic simulation of pilus rod assembly and assembly termination, Related to Figure 3C. Schematic of part of the pilus rod assembly mechanism used for Monte Carlo simulations in MATLAB and SimBiology. Each species occurring during the assembly reaction was represented by a dedicated pool. For example, species A, B and I denote activated FimDCH, FimCA and FimCI, respectively; species AB, AB2, AB3 and AB4 denote pili with 1, 2, 3 or 4 FimA monomers incorporated and ABI, AB2I and AB3I represent pili with 1, 2 and 3 FimA monomers that were terminated by incorporation of FimI. Dotted arrows indicate the direction of the continuing reaction.

**Figure S8.**
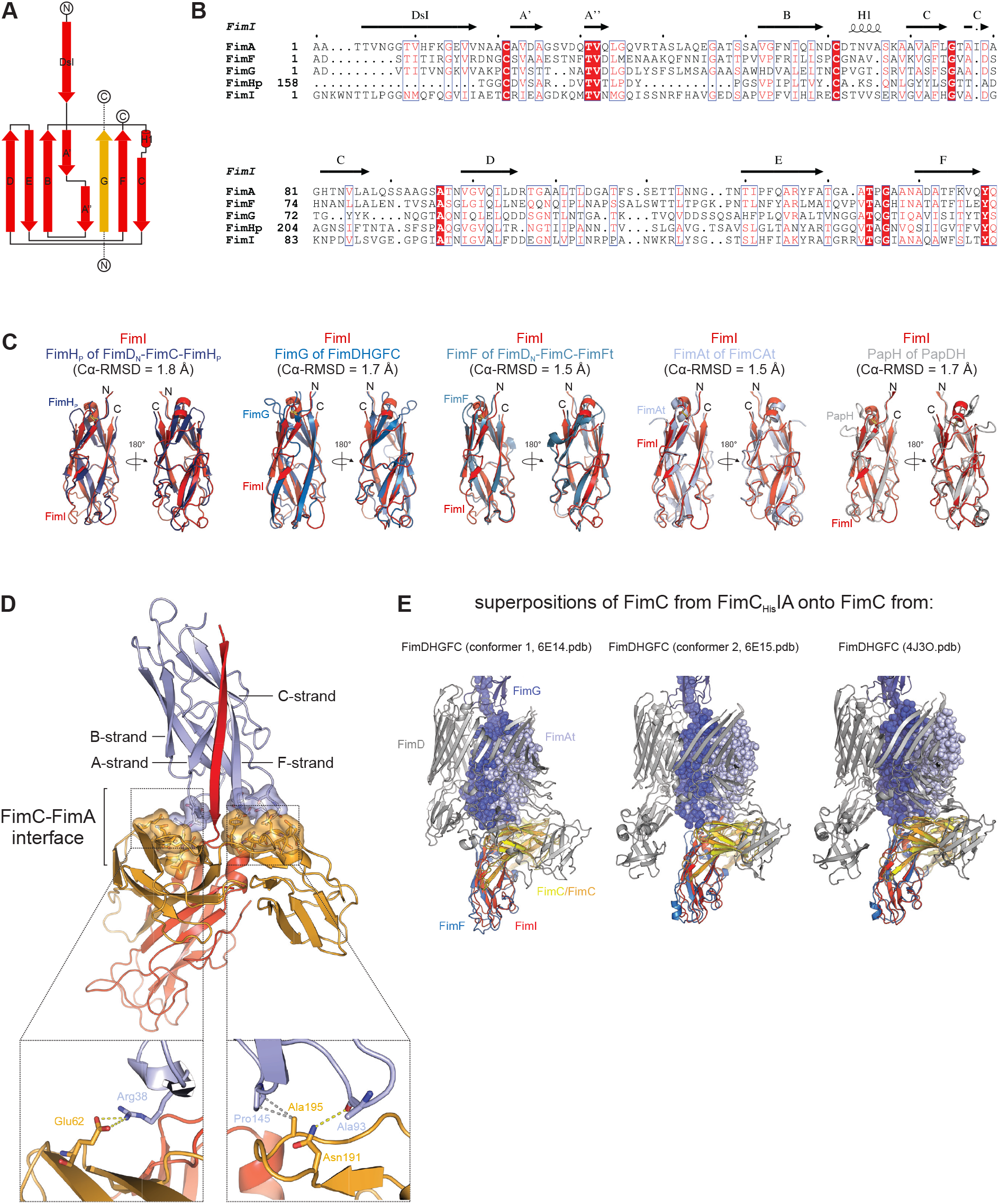
Structural analysis of the FimC_His_IA complex, Related to Figure 7. (A) Topology diagram of the FimI fold as found in the structure of the FimC_His_IA complex. β-strands are depicted as arrows, α-helices as cylinders. The FimC donor strand is shown in gold and runs parallel to the C-terminal F-strand of FimI. (B) Structure-based sequence alignment of all type 1 pilus subunits. Similar residues are in blue boxes and highlighted in red, identical residues are shown in white on red background. The secondary structure elements of FimI are displayed on top of the alignment. (C) Structural superposition of FimI of the FimC_His_IA complex onto FimH_P_, FimG, FimF, FimAt and PapH of the FimD_N_-FimC-FimH_P_ (1ZE3.pdb), FimDHGFC (4J3O.pdb), FimD_N_-FimC-FimF_t_ (3BWU.pdb), FimCAt (4DWH.pdb) and PapDH complex (2J2Z.pdb), respectively. (D) Crystal structure of the FimC_His_IA complex shown in cartoon representation with the 351 Å^2^ interface formed between FimC and FimAt shown in surface representation. FimC is in gold, FimI in red and FimAt in light blue. Details of the FimC-FimAt interaction are shown in boxes at the bottom of the panel. The salt bridge, hydrogen bond and hydrophobic interactions are indicated as dashed yellow and gray lines. (E) Structural superposition of FimC from the FimC_His_IA complex onto FimC from the FimDHGFC complex. All proteins are shown in cartoon representation except for FimG and FimAt, which are shown as spheres. FimD is in gray, FimG in blue, FimF in marine, FimC in yellow/gold, FimI in red and FimAt in light blue. Note that while FimF and FimI superimpose well, FimAt clashes with FimD and does not align with FimG inside the pore of FimD.

**Table S1.**
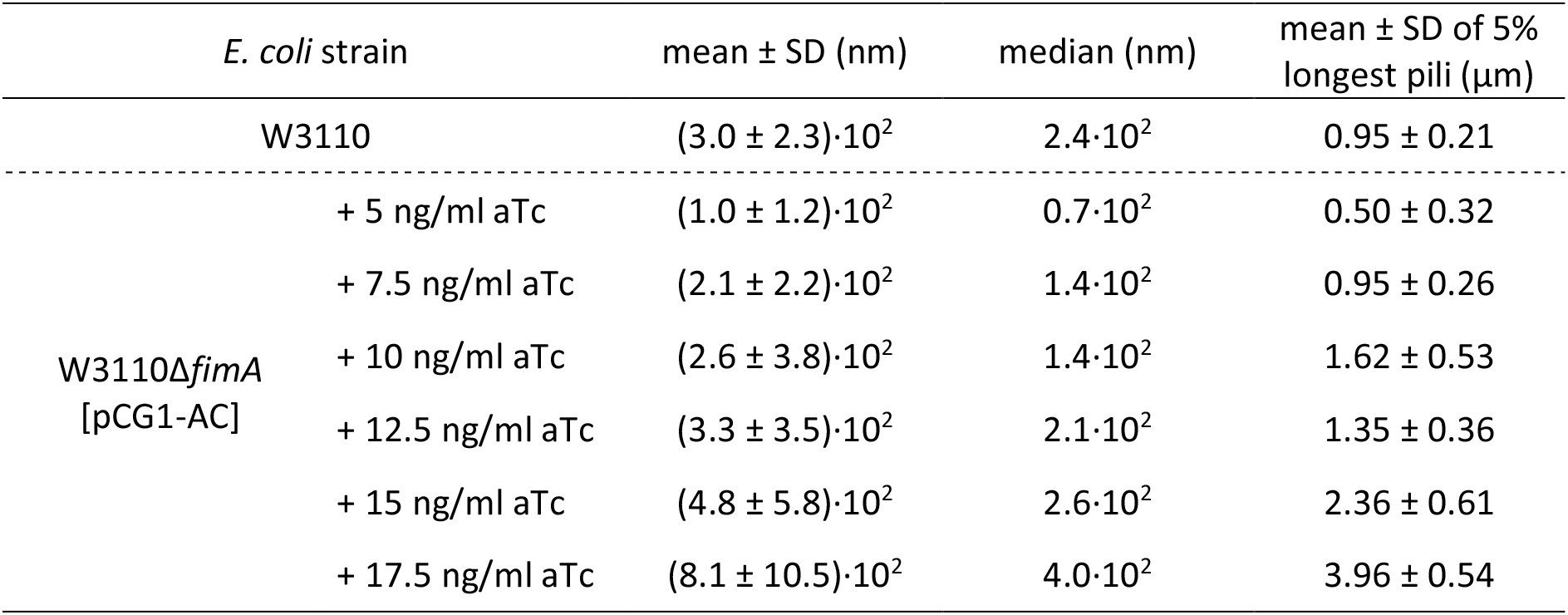
Parameters of pilus length distributions obtained after *in vivo* pilus assembly, Related to Figure 4C. Pili were released from *E. coli* W3110 or W3110Δ*fimA* [pCG1-AC] grown in presence of the indicated anhydrotetracycline (aTc) concentrations and parameters obtained from length measurements of 200 pili each. SD denotes the standard deviation of the mean.

**Table S2.**
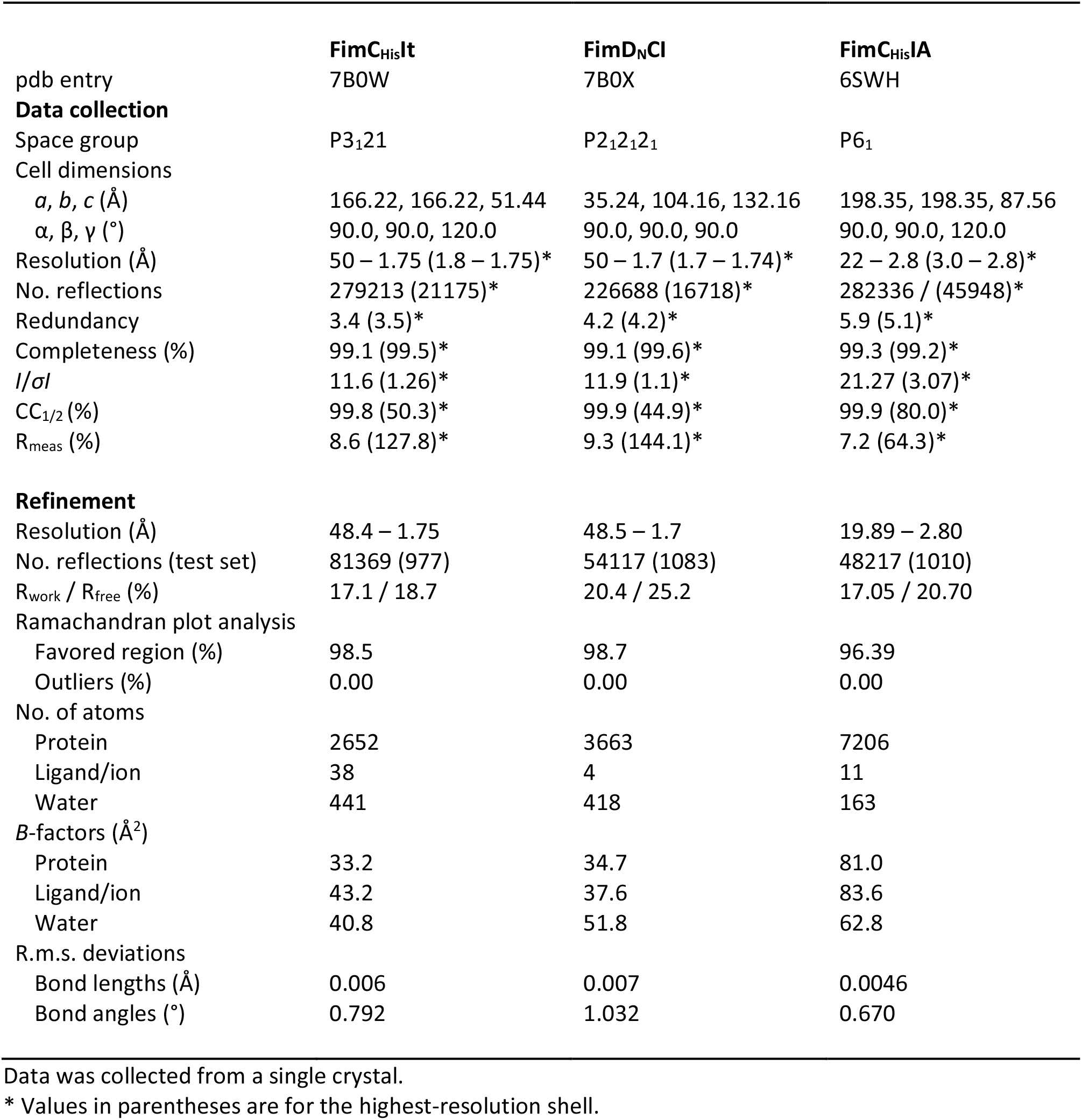
X-ray data collection and refinement statistics for structure determination of the FimC_His_It, FimD_N_CI and FimC_His_IA complexes.

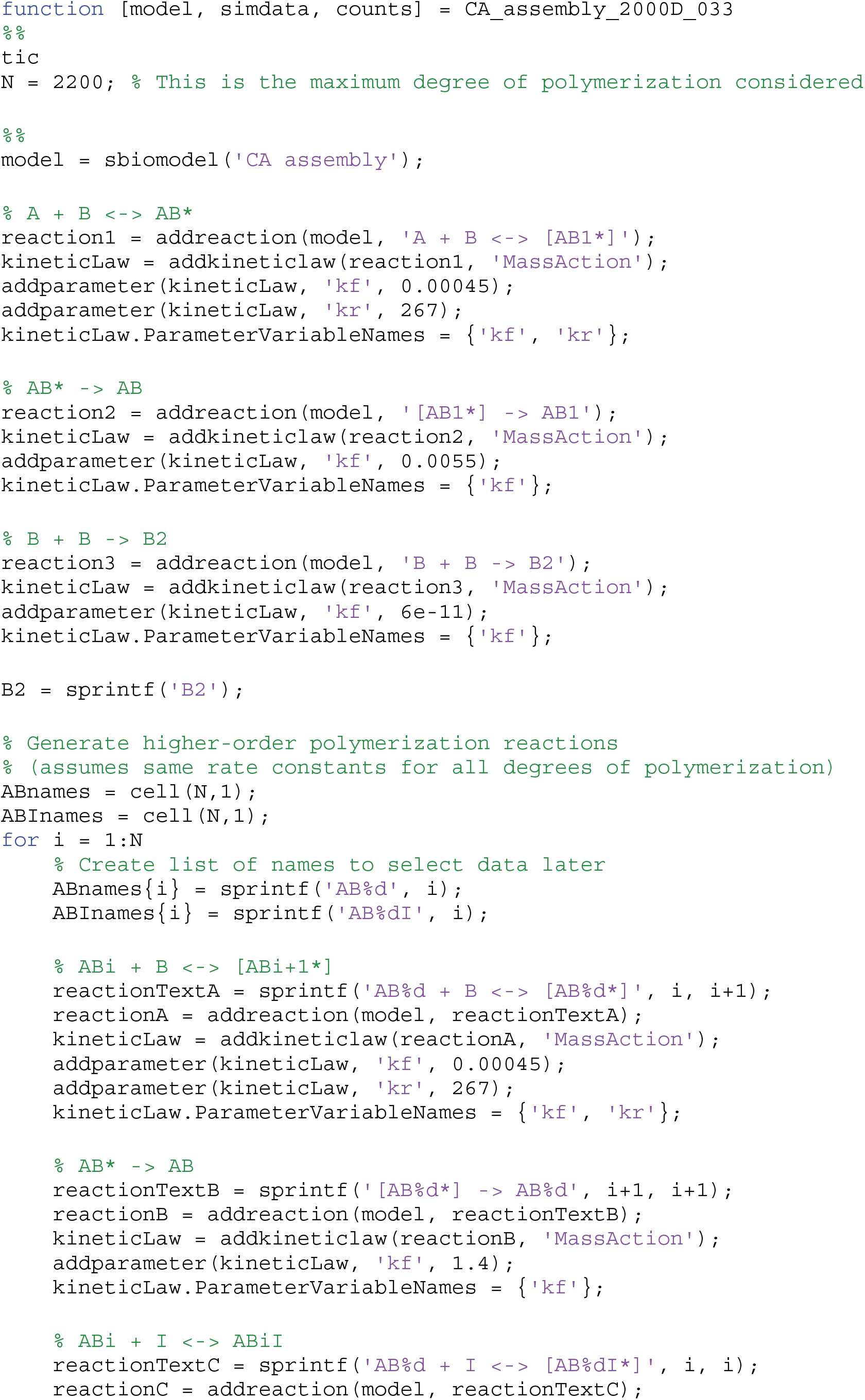
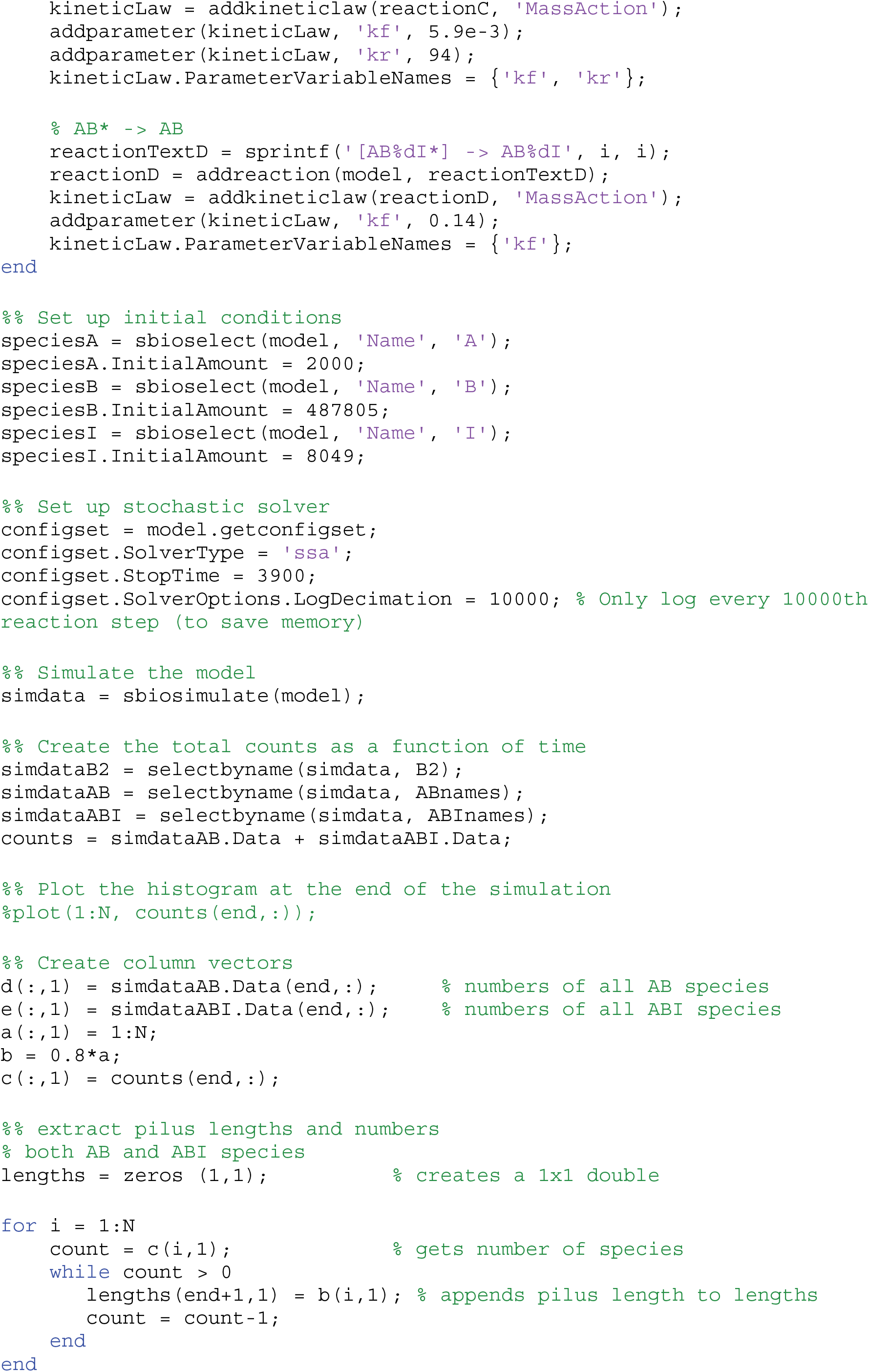
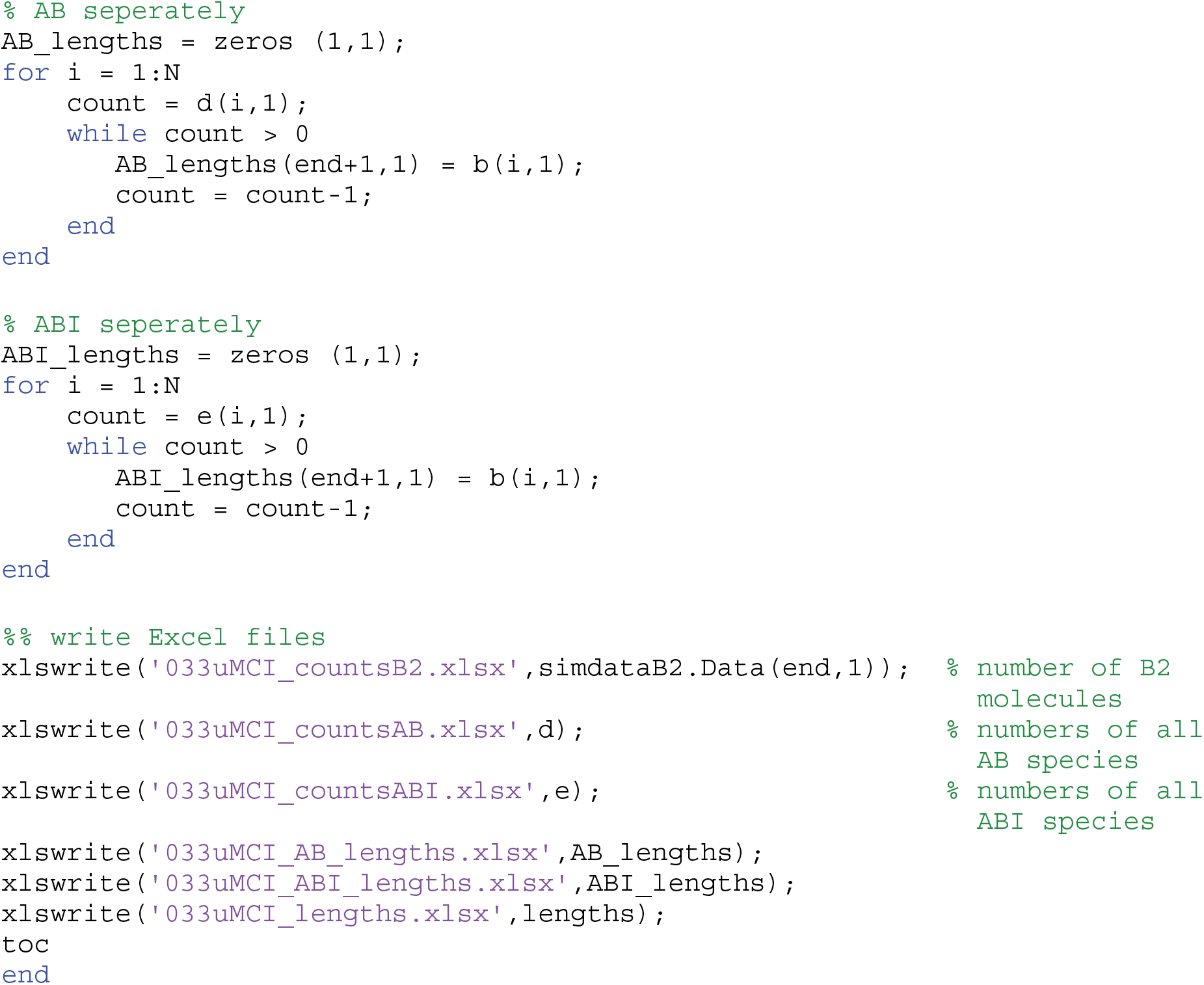
MATLAB script used for Monte Carlo simulations of pilus rod assembly reactions in presence or absence of FimCI.

